# Genetic basis of cuticular hydrocarbon variation in the desert ant *Cataglyphis niger*

**DOI:** 10.1101/2025.03.30.646157

**Authors:** Shani Inbar, Besan Saied, Pnina Cohen, Zeev Frenkel, Yoann Pellen, Abraham B. Korol, Eyal Privman

## Abstract

Cuticluar hydrocarbons (CHCs) are a ubiquitous component of insect cuticles that are used for a wide range of chemical signaling functions, especially recognition. Recognition and other signals are vital for the maintenance of insularity and cooperation in social insect colonies. Therefore, we expect natural selection on the composition and variabitlity of social insect CHC profiles. Selection on these signals may result in the evolution of genetic polymorphism affecting variation in CHC profiles. Here we tested for a genetic basis of CHC variation in the desert ant *Cataglyphis niger*. We applied a genomic mapping appraoch to a cohort of brothers from the same nest to reduce noise from environmental effects and achieve a clear statistical signal for association between the variation of CHCs and quantitative trait loci (QTL). This analysis identified 19 QTLs associated with 8 out of the 31 CHCs identified, and one QTL associated with total CHC quantity. These QTLs are located on 11 different chromosomes, including two cases where QTLs of different CHCs overlap. Each QTL explains between 13-25% of the variation in a specific CHC. We highlight several candidate genes in the QTLs identified, including fatty acid elongase and reductase genes. Our results reveal a polygenic genomic architecture underlying CHC variation in a population of the desert ant and open new research avenues into the genetic basis and evolution of chemical signaling in social insects.

## Introduction

Long-chain cuticular hydrocarbons (CHCs) are a major component of the cuticular lipids of insects, which function as touch pheromones communicating a range of chemical signals (Drijfhout, et al. 2009). For example, males and females display distinct, species-specific CHCs and other cuticular lipids that are used for mate choice in different species, including *Drosophila* fruit flies (Ferveur and Cobb 2010; Jezovit, et al. 2017; Khallaf, et al. 2021) and *Nasonia* parasitoid wasps (Steiner, et al. 2006; Niehuis, et al. 2013; Buellesbach, et al. 2022). In social insects, CHCs are also used for nestmate recognition. The ability to discriminate nestmates from foreign intruders is essential for colony integrity and protection of colony resources. CHCs are the main chemical cues underlying nestmate recognition (Holldobler and Michener 1980; Singer 1998). In this study, we investigated the genetic basis of CHC variability in terms of the number of loci and their effect sizes and identified candidate genes that may affect this important phenotype.

CHCs are the main lipid constituents of the epicuiticular layer of insects (reviewed in Blomquist and Bagneres 2010; Blomquist, et al. 2020). It has been speculated that the ancestral role of CHCs was probably to provide a waterproof layer preventing desiccation (Hadley 1994; Blomquist and Bagnères 2010 p. 5). Subsequently, they evolved a secondary role as chemical cues for signaling, communication, and recognition. Insect CHCs have variable chain lengths (20 to 50 carbons) and variable number and position of double bonds and methyl branches, thus allowing a great diversity that affords high information content as chemical cues. CHCs function as sex pheromones in many insects, as attractants, as fertility signals, and as chemical cues for species recognition that facilitates mate choice (Pickett, et al. 2009; Ferveur and Cobb 2010). Different social insect castes generally display distinct CHC profiles, including distinction between fertile individuals (queens, gamergates, or egg laying workers) and their less fertile or non-reproductive nestmates (Howard, et al. 1982; Nunes, et al. 2009; Kleeberg, et al. 2017; Derstine, et al. 2018; Sprenger and Menzel 2020). Signaling by CHCs have been extensively studied in social insects, where they make up an important component of the elaborate communication system necessary for coordinating complex collective behaviors (Hefetz 2007).

The enzymatic pathways responsible for the biosynthesis of CHCs were not characterized in social insects, however, a large body of literature describes such enzymes in model insect species, most notably *Drosophila melanogaster* (Blomquist 2010; Holze, et al. 2021). CHCs are *de novo* synthesized from two- or three-carbon units. Short saturated fatty acids (up to 18 carbons) are first synthesized by fatty acid synthases. These precursors are then elaborated by a series of enzymes that elongate the carbon chain (elongases), introduce double bonds (desaturases), and the final enzymatic steps that convert the fatty acid to a hydrocarbon, including reductases and decarboxylases. Hymenopteran homologs of these enzymes have been identified and studied for their role in CHC biosynthesis in bees and wasps (e.g., Falcón, et al. 2014; Buellesbach, et al. 2022; Moris, et al. 2023). Homologs are also found in ant genomes, and are the natural candidates for the genetic basis of ant CHCs (e.g., Sprenger, et al. 2021; Wen, et al. 2025; and our own observations). Some of these gene families have significantly expended in ants, indicating the evolution of greater complexity in CHC synthesis (e.g., Hartke, et al. 2019). Beyond the genes involved in synthesis of CHCs, other genes may be involved in regulation of the quantitates of CHCs produced and secreted through the cuticle. These genes may function in any stage of the long road from the site of CHC production to the external surface of the cuticle. CHCs can be either synthesized de novo or from diet-derived fatty acids (Piek 1964; Blomquist and Jackson 1973; Liang and Silverman 2000; Blomquist 2010). They are thought to be produced primarily in cells known as oenocytes in the fat body, and then transported through the hemolymph, secreted and deposited on the cuticle (Gu, et al. 1995; Fan, et al. 2003).

One of the best understood social functions of CHCs is in nestmate recognition, a fundamental and vital function in social insect colonies. Nestmate recognition is primarily based on chemical recognition, which was suggested to involve matching an olfactory perception of chemical cues to an experience-derived neural template (Ozaki and Hefetz 2014). Different species display different mixtures of CHCs (Martin and Drijfhout 2009a; d’Ettorre and Lenoir 2010), which suggests rapid evolution, possibly driven by strong selection. Intra-specific quantitative variation generates colony-specific CHC profiles that facilitate nestmate recognition, as demonstrated by experimental manipulation in multiple ant species (Bonavita-Cougourdan, et al. 1987; Nowbahari, et al. 1990; Akino, et al. 2004; Ozaki, et al. 2005; Martin, et al. 2008), including in our study species *Cataglyphis niger* (Lahav, et al. 1999). These studies transferred the hydrocarbon fraction of extracts from one ant to another, which caused ants to attack nestmates or to reduce aggression towards non-nestmates. Some studies went a step further: they synthesized specific hydrocarbons and tested their effect on nestmate recognition (Akino, et al. 2004; Martin, et al. 2008).

Recognition cues may be heritable and/or environmentally derived. Heritability of CHC profiles would imply that quantitative variations in CHCs are partially explained by genetic polymorphism. Both genetic and environmental effects were documented in *Drosophila*, and some of the underlying genes were identified (reviewed in Ferveur 2005). Multiple studies suggest that quantitative variation in CHCs is heritable in diverse insect species including crickets (Thomas and Simmons 2008), termites (Dronnet, et al. 2006), and parasitoid wasps (Niehuis, et al. 2011; Buellesbach, et al. 2022). Empirical studies in social insects suggest that CHC variation is affected by both environmental and genetic factors: For example, a strong effect of the diet was reported in the Argentine ant (Liang and Silverman 2000).

Pathogens may also have an effect on nestmate recognition in ants (Tranter, et al. 2015; Leclerc and Detrain 2016). Studies in ants and bees reported the effect of parasites and pathogens on CHCs and nestmate recognition (Trabalon, et al. 2000; Beros, et al. 2017; Cappa, et al. 2019). Other potential factors may include abiotic effects such as temperature and humidity (Buellesbach, et al. 2018; Menzel, et al. 2018; Sprenger, et al. 2018; Silva, et al. 2024). Nevertheless, genetic effects may also be involved, and multiple studies suggested that ant CHCs are heritable to a certain degree: Several studies demonstrated that intercolonial variation in CHC profiles and the ability for nestmate recognition are retained under controlled conditions and uniform diet (Nowbahari, et al. 1990; Lahav, et al. 1999; Akino, et al. 2004; Ozaki, et al. 2005; van Zweden, et al. 2009). Behavioral experiments on different populations of the ant *Temnothorax longispinosus* revealed an association between aggression and genetic dissimilarity (Stuart and Herbers 2000). Isolation of *Formica exsecta* workers revealed a weak patriline effect on CHCs, indicating that different alleles inherited from different fathers had an effect on the CHC profiles of their daughters (Martin, et al. 2012). A study in the ant *Monomorium pharaonis* tested for heritability of individual CHCs by systematic intercrossing eight colonies over nine generations (Walsh, et al. 2020), resulting in low to moderate estimated heritability: narrow-sense heritability *h^2^* between 0.006 and 0.36. These studies provide evidence for heritability of CHCs in diverse ant species.

The vital function of nestmate recognition is expected to have large fitness consequences, including “red queen” dynamics due to selection against social parasites (Blomquist and Bagneres 2010). This hypothesis is supported by the great interspecific diversity of CHCs and their rapid evolution (Martin and Drijfhout 2009a; d’Ettorre and Lenoir 2010). Some studies found stronger differentiation between colonies by methyl-branched vs. straight alkanes (e.g., Dahbi, et al. 1996). The abundance of branched alkanes and unsaturated alkenes in ant CHCs, which provide greater information content as cues while compromising protection from desiccation, was suggested to be the result of selection for signaling functions (d’Ettorre and Lenoir 2010; Sturgis and Gordon 2012). The exceptional complexity of ant CHC profiles suggests selection for recognition accuracy. If genetic polymorphism arises such that it affects nestmate recognition cues, selection for distinctness of colony profiles is expected to increase the frequency of novel alleles. The requirement for distinctness of colony CHC profiles should lead to selection in favor of rare alleles, i.e. negative frequency-dependent selection resulting in balancing selection (Ayala and Campbell 1974; Jongepier and Foitzik 2016). Such processes result in a large number of alleles per locus and the maintenance of highly diverged alleles in the population over long evolutionary time, as observed for example in mammalian Major Histocompatibility Complex (MHC) genes (Hughes and Nei 1988) and the honeybee Complementary Sex Determination (CSD) locus (Cho, et al. 2006). We propose that the critical role of CHCs in nestmate recognition is expected to result in such evolutionary dynamics, which would lead to the accumulation of polymorphic loci that affect the display of ant CHCs quantitatively, i.e., quantitative trait loci (QTLs).

Here, we conducted a QTL mapping study to test whether quantitative variation of CHCs is associated with genetic polymorphism in the desert ant *Cataglyphis niger*. We produce the genomic infrastructures necessary for mapping associated genomic loci – a chromosome-level reference genome sequence. We conduct reduced representation genomic sequencing of a cohort of brothers to identify polymorphic loci associated with CHC variation (QTLs) and characterize the genomic architecture of these traits. The use of haploid males affords valuable advantages in terms of the simplicity of the underlying genetics and minimal environmental variation (see Methods). The QTLs we discovered in males are most likely to have a functional role in shaping their CHC profiles that are used as cues for mate choice, i.e., as sex pheromones. However, we propose that these QTLs are likely to also have effects on CHCs in workers and queens because many of the same CHCs are observed in all castes of the species (Hefetz 2007; Inbar and Privman 2019) and because the biosynthetic pathway of a given CHC is likely to be similar in all castes. We highlight promising candidate genes in these loci. The discovery of genetic loci associated with CHC variation serves as a first step in the study of the molecular mechanisms underlying chemical signaling functions including sex pheromones and potentially also nestmate recognition, and their evolution in a social insect species.

## Methods

### Samples

Our analysis was based on a sample of 117 brothers, collected on 3/5/2016 from a single *C. niger* nest in Betzet Beach in the northern Israeli coastline (colony BZT50, 33°03’32N 35°06’15E) and frozen in −80°C. Each of these samples were used for chemical analysis and genomic sequencing. This colony belongs to a population that was previously referred to as *Cataglyphis drusus* (Eyer, et al. 2017), but our species delimitation analysis showed that it is part of the species described as *C. niger* that is widespread across Israel (Brodetzki, et al. 2019). Our population structure analysis of nuclear genomic DNA sequence data revealed no sub-structure of genetically distinct population clusters –populations across Israel are not genetically differentiated (Brodetzki, et al. 2019). We note that in some areas in Israel *C. niger* colonies have multiple queens (polygyne) and that our recent study revealed that this social polymorphism is associated with a supergene on chromosome 2 (Lajmi, et al. 2025). However, the study population used here is purely monogyne, i.e., each colony is headed by a single queen (Brodetzki, et al. 2019). The supergene is always homozygous (MM) in monogyne colonies (the P haplotypes is only found in polygyne colonies), so we do not expect it to affect the QTL mapping analysis in this study.

Using a cohort of brothers for QTL mapping affords several advantages. Ants have haplodiploid sex determination, where haploid males develop from unfertilized eggs, and diploid females derive from fertilized eggs. Thus, males have no fathers. They only receive one of their mother’s alleles in each locus. And so, a cohort of brothers resents the products of maternal meiosis. Such a cohort servses as a simple mapping population similar to a back-cross experiment that is traditionally used for QTL mapping studies. Moreover, recessive alleles are more readily revealed in haploid progeny because samples are hemizygous at all loci, whereas recessive effects might not be observable in diploid samples (in heterozygous samples). For the same reason, similar studies using haploid males were conducted in another hymenopteran species – the parasitoid wasp *Nasonia vitripennis* (Niehuis, et al. 2011). This approach successfully identified a large number of QTLs associated with many of the CHCs analyzed. Another advantage is that CHC variation among brothers sampled from the same nest is likely to be minimally affected by environmental factors because they all experienced a similar environment in the nest. For example, we can safely assume they were fed the same diet and exposed to the same microbes by the workers that cared for them. Males are also expected to be more uniform than workers because they do not have distinct task groups that are exposed to different environments (e.g. nurses vs. foragers).

### Chemical analysis

After being kept in a −80 freezer, a random substet of 81 of the 117 samples were individually extracted in 100 μl hexane, containing 200 ng tetracosane (C24) as internal standard.

Compounds were identified by combined gas chromatography/mass spectrometry (GC/MS) based on their fragmentation patterns, as compared to authentic standards (linear aliphatic hydrocarbons from C10 to C40 supplied by Supelco), following the methods used in previous studies of this and other populations of the species (e.g., Brodetzki, et al. 2019). GC/MS was conducted using an Agilent 19091S-433UI instrument at EI and splitless modes, equipped with an HP-5ms (Ultra Inert) column, temperature-programmed from 60°C to 300°C (with 1 min initial hold) at a rate of 5°C per min, with a final hold of 15 min. Each sample was then quantified by gas chromatography. Peak integration was performed using the program Varian Galaxie version 1.9 to obtain the relative quantities of CHCs. We chose the conservative approach of using a baseline that connects the vallies on either side of a peak in order to avoid overestimation due to the effect of large peaks on neihboring smaller peaks. Conversely, this approach might result in underestimation. Absolute quantities were calculated relative to the internal standard, however the QTL mapping analysis was done using the relative quantities.

### Reference genome sequencing, assembly and annotation

Identification of the genes inside QTLs requires a genome assembly of sufficient contiguity, that is, genomic scaffolds should be large enough so that the markers of the linkage map can be linked to the genes within the confidence interval for the position of a QTL. We already had a reference genome assembly for *C. niger* that was assembled from Illumina short-read sequencing of a single haploid male (version Cnig_gn1; Yahav and Privman 2019). However, this assembly was rather fragmented, with a scaffold N50 size of only 17.9 Kb. Therefore, we conducted Pacbio long-read sequencing to improve the assembly contiguity. High-molecular weight DNA was extracted from a pool of ∼30 pupae from a *C. niger* colony also collected from the Betzet Beach population. A library was constructed and sequenced (CLR sequencing) on a Pacbio Sequel II machine at the Roy J. Carver Biotechnology Center, University of Illinois at Urbana-Champaign. The sequencing yielded 7,031,744 reads, with 169 Gbp in total, and an average subread length of 7,227 bp.

The Pacbio data was assembled into two versions of the genome, Cnig_gn3.1 and Cnig_gn3a.1. Version Cnig_gn3.1 was built more stringently, aiming to include only confidently assembled contigs, while version Cnig_gn3a.1 aimed to map as much of the genome sequence to chromosomes. We note that some artefacts are likely in the latter version, which may be harmful to various types of analyses. We only use this assembly for identifying candidate genes in QTLs. For this purpose, this practice is unlikely to have serious detrimental effect. To avoid technical artifacts in Pacbio sequencing process short (<5 kbp) and long (>50 kbp) reads were excluded from assembly Cnig_gn3.1. Reads were assembled onto contigs using *Canu* version 2.1.1 (Koren, et al. 2017). Contigs of Cnig_gn3a.1 were based on all of the PacBio reads. Contigs of both assembly versions were polished by *Pilon* version 1.24 (Walker, et al. 2014) using the short Illumina reads from assembly Cnig_gn1. Polished contigs were anchored to the linkage map (see below) based on markers with a unique *blastn* hit to contigs. Contigs with no mapped markers were fitted into chromosomes based on the local similarity between the genomes of *C. niger* and *Formica selysi* (https://www.ncbi.nlm.nih.gov/assembly/GCA_009859135.1/) if the synteny of the two genomes was colinear in the relevant chromosomal region based on *blast* hits of *C. niger* genetic markers to *F. selysi* chromosomes and *blast* of *F. selysi* genes to *C. niger* contigs. Chromosomes in the Cnig_gn3.1 assembly contain only contigs with unique position and defined orientation. Cnig_gn3a.1 also includes contigs with non-unique mapping and/or unknown orientation. Contigs with several (up to 5) possible positions were included in Cnig_gn3a.1 several times. Therefore, the Cnig_gn3a.1 had greater total length (265 vs. 236 Mbp) and N50 contig size (3.8 vs. 1.1 Mbp) relative to Cnig_gn3.1. The Cnig_gn3a.1 assembly includes 83% of the total assembly length in chromosomes (i.e. 17% of the sequence is in contigs that no mapped markers), whereas the Cnig_gn3.1 assembly includes 74%.

Gene annotation was conducted using the *GeMoMa* pipeline (Keilwagen, et al. 2016; Keilwagen, et al. 2018) that incorporates three types of evidence: (1) RNA sequencing of samples from all life stages and castes (NCBI accession PRJNA494690; Yahav and Privman 2019) aligned to the genome using *HISAT2* (Kim, et al. 2019); (2) homology to genes annotated in other insect genomes (*Camponotus floridanus, Harpegnathos saltator, Linepithema humile, Solenopsis invicta, Pogonomyrmex barbatus, Formica exsecta, Apis mellifera, Nasonia vitripennis, Anopheles gambiae, Drosophila melanogaster, Tribolium castaneum*); and (3) *ab initio* gene prediction using *Augustus* (Stanke, et al. 2006). The annotation resulted in 14,889 and 16,260 putative protein-coding genes for Cnig_gn3.1 and Cnig_gn3a.1, respectively. Functional annotation of genes was assigned based on homology using the *Trinotate* tool (version 3.0.2; https://github.com/Trinotate/Trinotate.github.io/wiki) from the *Trinity* pipeline (Grabherr, et al. 2011), which uses *blast* against the Swissprot protein database and assigns genes to functional categories of the Gene Ontology (GO) database (Harris, et al. 2004). Out of the 16,260 genes in Cnig_gn3a.1, 10,268 genes had Swissprot hits, and GO terms were assigned 10,186 genes. To identify candidate genes in QTLs, we searched both Swissprot gene description and GO terms for keywords related to molecular functions and biological processes implicated in CHC synthesis, including fatty acid metabolism (keyword “fatty”) and development and function of the cuticle (keyword “cuticle”).

### RAD sequencing

DNA was extracted individually from each sample using TRIzol® (Invitrogen Life Technologies) and kept at −20°C. Quantity and quality of extracts were determined by NanoDrop™ 2000 Spectrophotometer and run on a 1.5% agarose gel to verify consistency in DNA yield, degradation, and phenol/salts residues. Reduced representation genomic libraries were constructed according to a modified double-digest Restriction-site Associated DNA sequencing (ddRAD-seq) protocol based on Brelsford, et al. (2016). Briefly, DNA was digested by two different restriction enzymes (EcoR1 rare-cutter; Mse1 frequent-cutter; from New England Biolabs) and ligated to barcoded adaptors for multiplexing (using custom oligos with 96 barcodes of variable lengths of 4-8bp, paired with two different barcoded PCR primers to allow multiplexing all 117 samples together). Amplified products were pooled and purified (Beckman Coulter, Agencourt® AMPure® XP) so that 200-700bp fragments would be selected. The library was paired-end sequenced on one lane of a HiSeq4000 Illumina sequencer.

The sequencing resulted in 693,969,414 150b reads (346,984,707 pairs), of which 662,096,312 were associated with a sample (barcode). The reads were quality checked and those in which the quality score drops below an average phred score of 10 (i.e. 1 in 10 chance for sequencing error) in a sliding window of 22 bases were removed using *process_radtags* from the *STACKS* pipeline (Catchen, et al. 2011). After this initial filtering a total of 630,112,110 reads remain, which is expected to give an average sequencing depth of ∼20X (based on 67,687 positions of the EcoRI restriction sites in the reference genome).

Reads were aligned to the reference genome using *Bowtie2* (Langmead and Salzberg 2012). Aligned reads that have more than 2 mismatches to the reference genome and reads that had multiple mappings were removed. The number of aligned reads retaind totaled 431,321,204, 62% of the original number of reads.

The aligned reads were analyzed together to create a catalogue of 88,439 SNPs using the *STACKS* pipeline, with an average sequencing depth of 14X across SNPs and samples. These SNPs were filtered using custom perl scripts to remove loci that had more than three samples had a heterozygous genotype call (not expected for haploid samples) or less than four samples with the rare allele (the brothers are expected to have roughly 50% of each of the maternal alleles) or more than 50% missing data. 7,769 SNPs passed this filtering.

### Linkage map

The program *MultiPoint ultra-dense* (Ronin, et al. 2017) was used to construct a linkage map, based on the genotypes of 7769 SNPs in the 117 samples. These loci were further filtered by *MultiPoint* for segregation distortion (based on Chi^2 test to reject the null hypothesis of 1:1 segregation with *p* < 0.005) and for missing data (no more than 23 missing genotypes), leaving 5466 SNPs. Clustering of linkage groups resulted in 27 clusters of SNP markers using a recombination frequency threshold of 0.28. Skeleton markers were selected in every linkage group (giving priority to “twin markers”) and ordered by an “evolutionary strategy” heuristic optimization, as implemented in *MultiPoint*. Ordering was assessed by jackknife resampling. We manually inspected the linkage groups and applied methods implemented in *MultiPoint* to merge linkage groups, exclude markers violating local map monotonicity, and complement the map by adding suitable singleton markers, resulting in a final map consisting of 26 linkage groups. To verify that the map length is not inflated by marker ordering errors, we checked the order by jackknifing and observed a very nearly perfect support across the map (i.e., 100% jackknife repeats gave identical order among each marker neighborhood along the linkage group length).

### QTL mapping

The program *MultiQTL* (Korol, et al. 1995) was used to map 32 quantitative traits of 81 samples, including 31 specific CHC traits and the total CHC amount. We defined the 31 specific CHC traits as relative amounts, while total CHC is an absolute amount in ng. Many of the traits had skewed, non-normal distributions, so we removed outliers and applied log-transformation where necessary. We first identified QTLs using the SIM (Single Interval Mapping) model in *MultiQTL*, which calculates a LOD score for every trait on each chromosome. For every putative QTL with LOD score greater than 1.8, we conducted a permutation test to obtain a *p*-value for the significance of association between genotypes and phenotypes. QTLs with *p*-value < 0.1 were carried on to the next step. For traits with two peaks of the LOD scores on a chromosome, we used the linked-QTL model to test the hypothesis of having two QTLs on the same chromosome (likelihood ratio test). For traits that had candidate QTLs on more than one chromosome, we applied the MIM (multi-interval mapping) model to all chromosomes identified in the SIM analysis, which generally results in higher LOD scores. The permutation test was repeated for every QTL using the MIM model. The SIM, linked-QTL, or MIM models also provided an estimate of the PEV (percent explained variation) for each QTL. We identified the location of each QTL by the peak of the LOD score, and its 95% confidence interval was defined as the region around the maximum, where the LOD score is greater than the maximum score minus 1.5 points. We applied the false discovery rate (FDR) method implemented in R (Benjamini and Hochberg 1995; Team 2021) to correct for multiple testing (32 traits by 26 chromosomes = 832 tests). We mapped the genetic position of the confidence interval of each QTL location to the genome assembly relative to the physical positions of genetic map markers. We then listed the annotated genes that reside within the scaffold regions contained in each QTL interval in assembly Cnig_gn3a.1.

## Results

### Reference genome and linkage map

As a basis for QTL mapping, we generated a new genome assembly for *C. niger* using high-coverage Pacbio sequencing that yielded improved contiguity compared to the first draft (scaffold N50 size of 1.08Mb compared to 17.9Kb for the first draft). We then constructed a linkage map using the genotypes of a cohort of 117 haploid brothers (Supp. File S1). We used RAD sequencing to genotype 7769 SNP loci that clustered into 26 linkage groups. We anchored the assembly contigs to the linkage map, yielding 26 chromosomal sequences that together encompass 83% of the total assembly length. The total map length was 5041cM, which is exceptional for animal species. Some of the chromosomes have a very long genetic map length: chromosome 1 is 402cM long and five additional chromosomes are longer than 280cM (Fig. 1). Given an estimated physical genome size of 236Mb (the total assembly length), we estimated the genome-wide average recombination rate at 21.3 cM/Mb.

**Figure 1:**
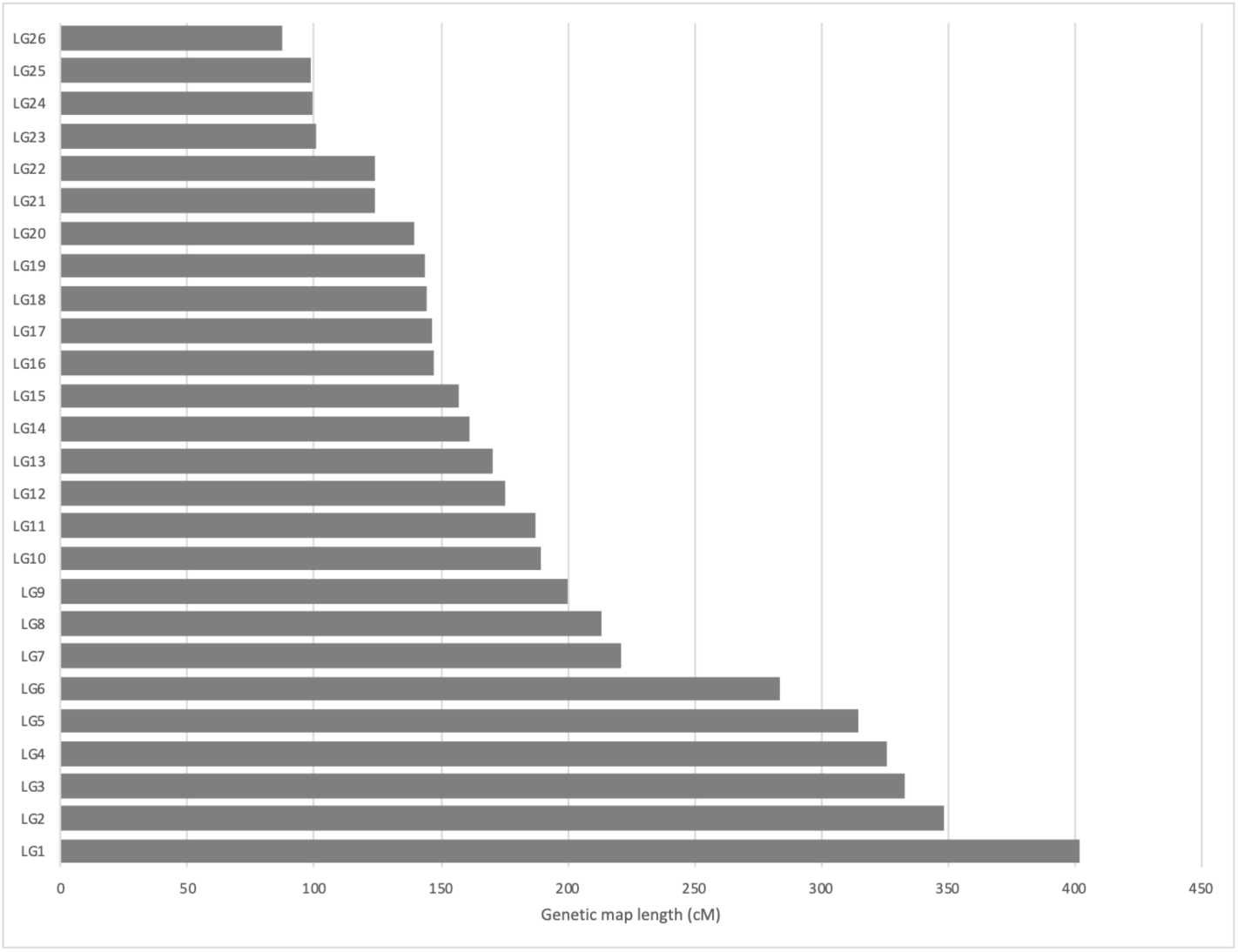
Distribution of genetic map lengths among the 26 chromosomes of *C. niger*.

### Chemical analysis of CHCs

We analyzed the cuticular extracts of the same brothers used for linkage map construction, which were treated as quantitative traits for QTL mapping. Chemical analysis identified 32 differentiated peaks in the chromatogram (Fig. 2; Table 1; raw data are provided in Supp. File S2). We were able to identify 36 CHCs in 32 peaks (four peaks consisted of two CHCs that failed to separate). These were saturated alkanes with carbon chain length ranging from C25 to C31, many of them having a methyl group, and some with two and even three methyl branches. We did not observe any unsaturated alkenes. Note that this means that our list of candidate genes for CHC synthesis will not include fatty acid desaturases, which are known from the biosynthetic pathway of alkenes in other insects. The quantitative traits for QTL analysis were defined as the proportion of each CHC out of the total CHC mixture. We also included the total CHC amount as an additional trait, which varied from 228 to 4825ng; average and standard deviation 1478±762ng. There was extensive variation both among and within traits. The most abundant peak contained 11Me- and 13Me-C29, and the second most abundant was C29. These two peaks had averages and standard deviations of 226±122ng and 168±121ng, respectively. Conversely, the least abundant CHC, 5Me-C27, was only 1.1±1.8ng, and undetectable in 34 samples. Generally, the distributions of all CHCs were asymmetrical, with many low values and a long right-hand tail, which required transformation and removal of outliers before QTL analysis (see Methods and Supp. Fig. S1).

**Figure 2:**
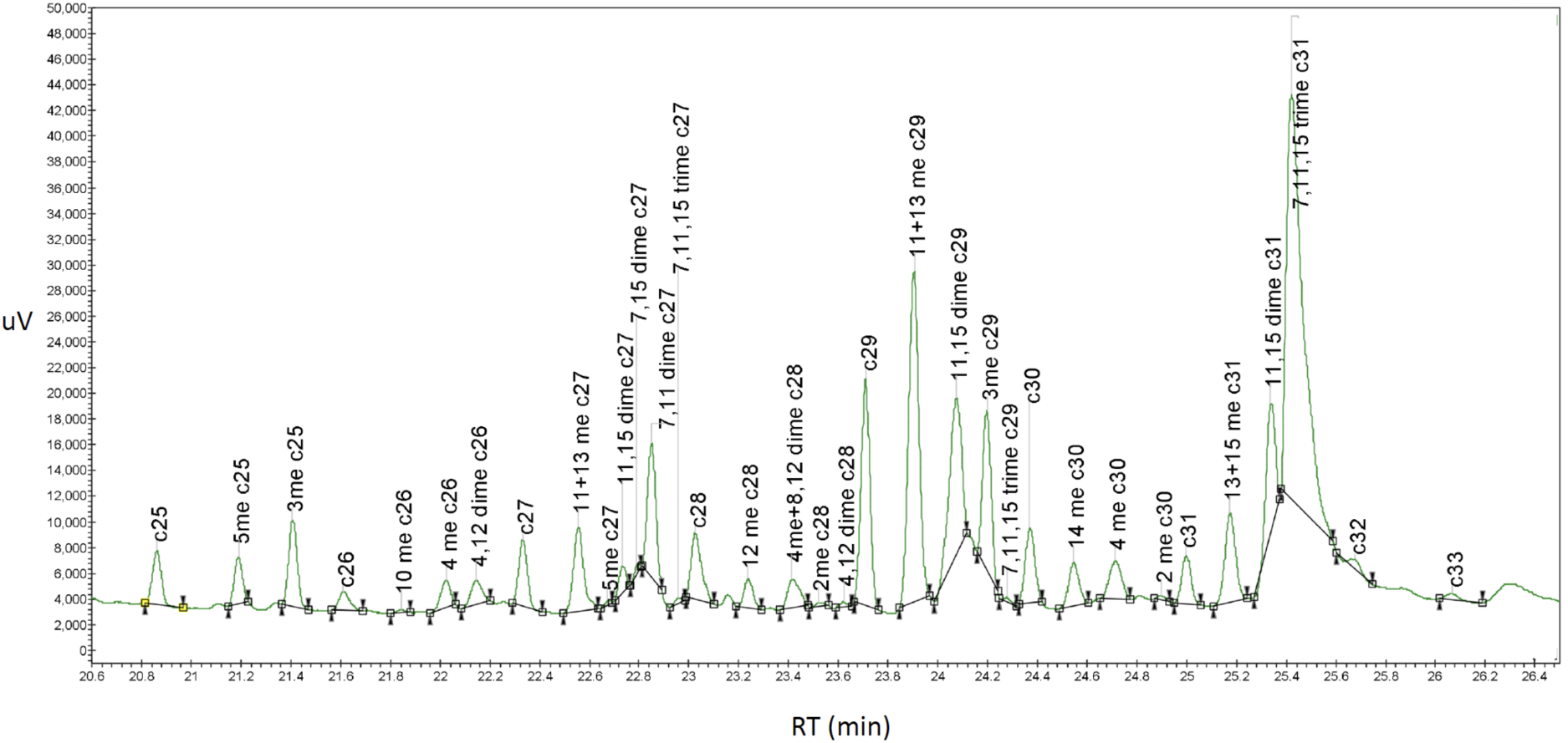
A chromatogram (uV absortion vs. retention time; RT) of the non-polar fraction (RT 20.5-26.5) of cuticular extracts from a *C. niger* male. Marked peaks were annotated by mass spectrometry.

**Table 1:**
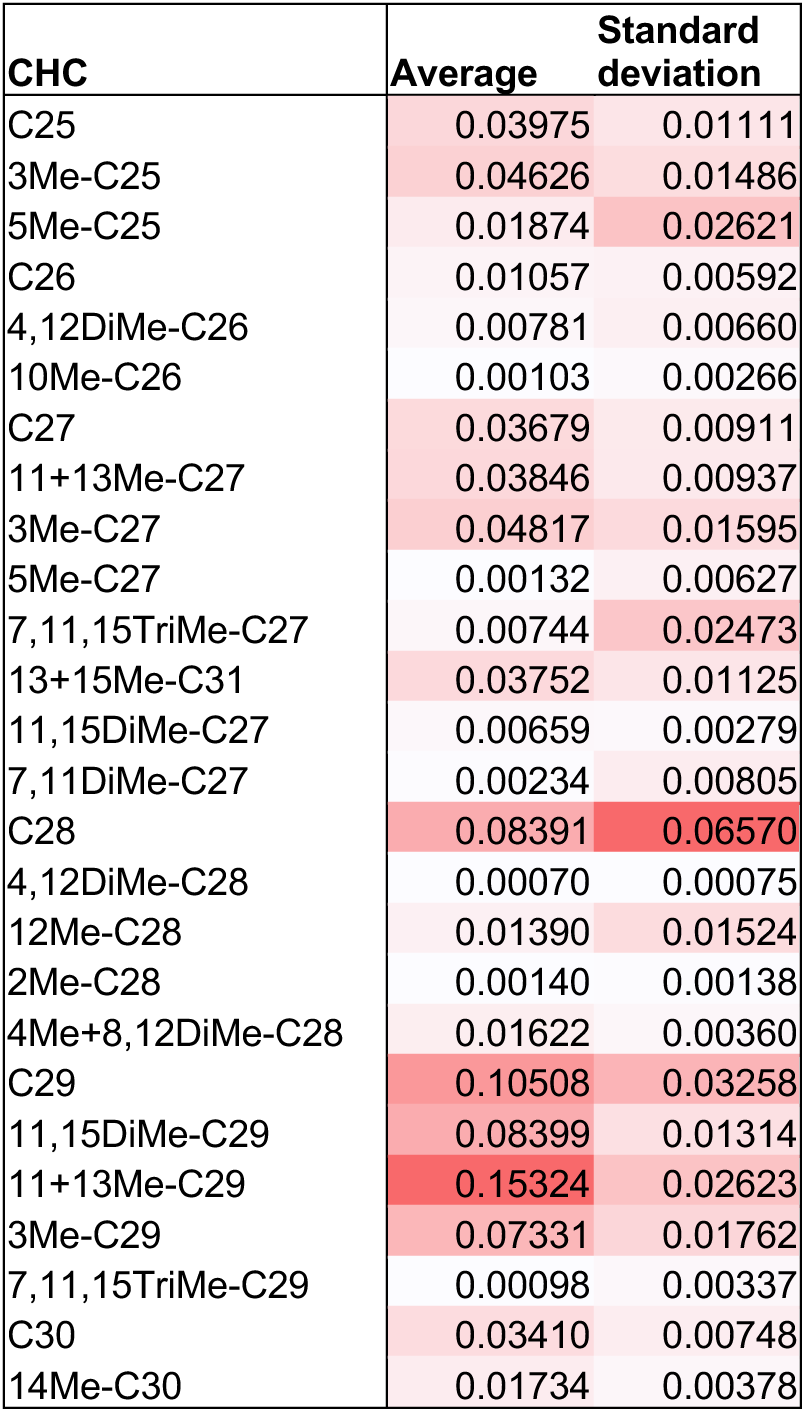
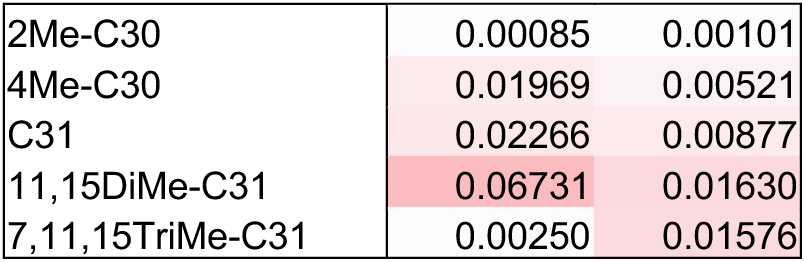
CHCs detected and their relative quantities in the cuticular extracts of *C. niger* males.

| CHC | Average | Standard deviation |
| --- | --- | --- |
| C25 | 0.03975 | 0.01111 |
| 3Me-C25 | 0.04626 | 0.01486 |
| 5Me-C25 | 0.01874 | 0.02621 |
| C26 | 0.01057 | 0.00592 |
| 4,12DiMe-C26 | 0.00781 | 0.00660 |
| 10Me-C26 | 0.00103 | 0.00266 |
| C27 | 0.03679 | 0.00911 |
| 11+13Me-C27 | 0.03846 | 0.00937 |
| 3Me-C27 | 0.04817 | 0.01595 |
| 5Me-C27 | 0.00132 | 0.00627 |
| 7,11,15TriMe-C27 | 0.00744 | 0.02473 |
| 13+15Me-C31 | 0.03752 | 0.01125 |
| 11,15DiMe-C27 | 0.00659 | 0.00279 |
| 7,11DiMe-C27 | 0.00234 | 0.00805 |
| C28 | 0.08391 | 0.06570 |
| 4,12DiMe-C28 | 0.00070 | 0.00075 |
| 12Me-C28 | 0.01390 | 0.01524 |
| 2Me-C28 | 0.00140 | 0.00138 |
| 4Me+8,12DiMe-C28 | 0.01622 | 0.00360 |
| C29 | 0.10508 | 0.03258 |
| 11,15DiMe-C29 | 0.08399 | 0.01314 |
| 11+13Me-C29 | 0.15324 | 0.02623 |
| 3Me-C29 | 0.07331 | 0.01762 |
| 7,11,15TriMe-C29 | 0.00098 | 0.00337 |
| C30 | 0.03410 | 0.00748 |
| 14Me-C30 | 0.01734 | 0.00378 |
| 2Me-C30 | 0.00085 | 0.00101 |
| 4Me-C30 | 0.01969 | 0.00521 |
| C31 | 0.02266 | 0.00877 |
| 11,15DiMe-C31 | 0.06731 | 0.01630 |
| 7,11,15TriMe-C31 | 0.00250 | 0.01576 |

One of these peaks (7,11,15TriMe-C31) was excluded from the analysis because its distribution was extremely non-normal, even after log transformation. Finally, 31 CHC traits were carried forward to the QTL mapping analysis.

### QTLs associated with variation in CHCs

Each CHC (a peak in the GC chromatograms) was treated as a quantitative trait and tested for association with the map markers (SNPs) in a QTL mapping analysis. The analysis included the relative amounts of 31 CHCs, and the absolute total amount of CHCs as an additional trait. A preliminary analysis identified 62 candidate genomic intervals with LOD scores greater than 1.8. Each candidate was evaluated by permutation analysis, which yielded *p*-values that were then corrected for multiple testing to control the false discovery rate (FDR). This analysis resulted in 20 QTLs (FDR < 10%) for nine traits, including one QTL for total CHC quantity (Table 2). The location and 95% confidence interval of each QTL was estimated based on the LOD function (Fig. 3).

**Figure 3:**
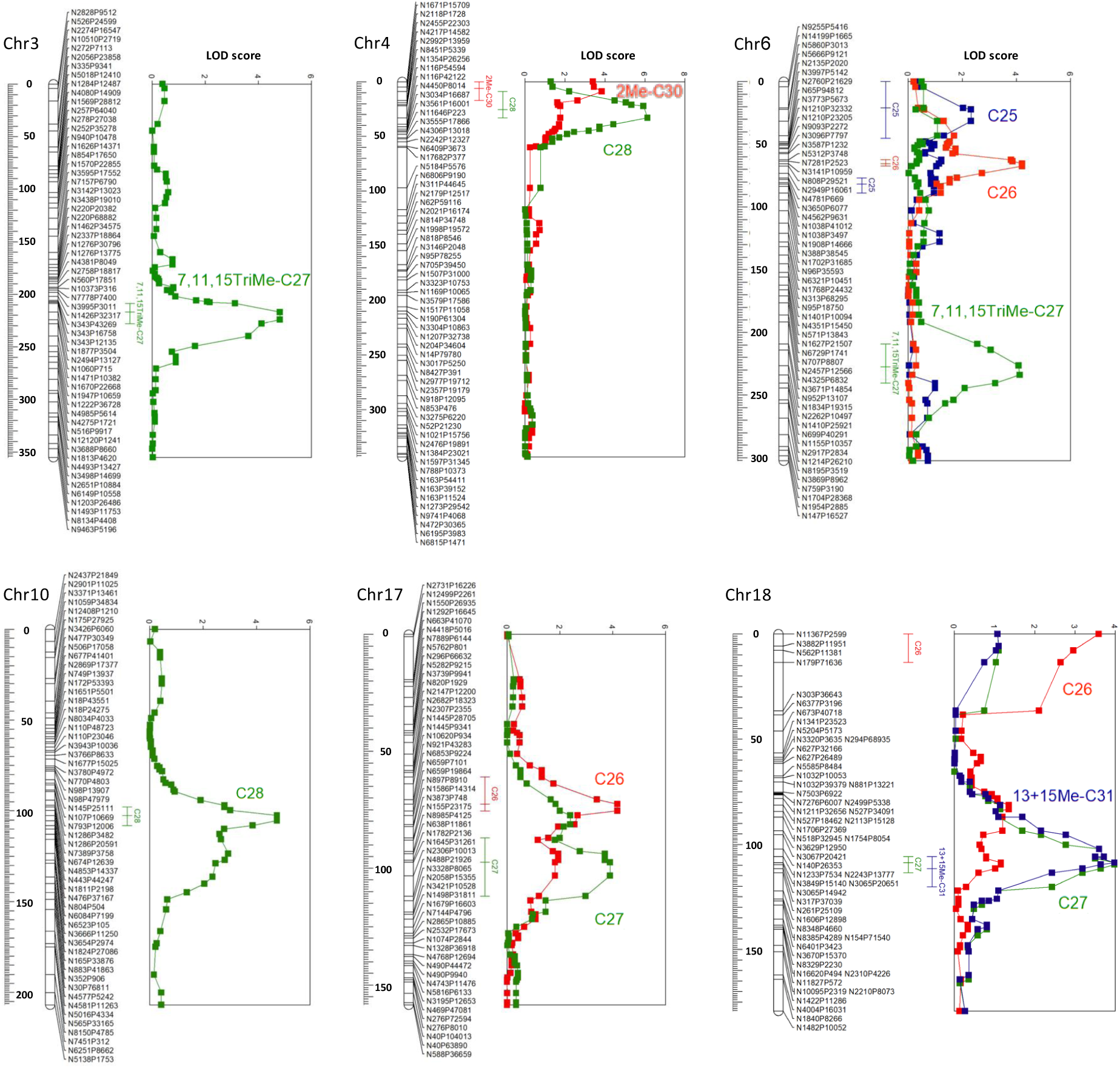
Location of CHC QTLs on six of the *C. niger* chromosomes. Map markers are listed with distances in cM along each linkage group. LOD scores are plotted for each trait, and 95% confidence intervals of QTL locations are indicated.

**Table 2:**
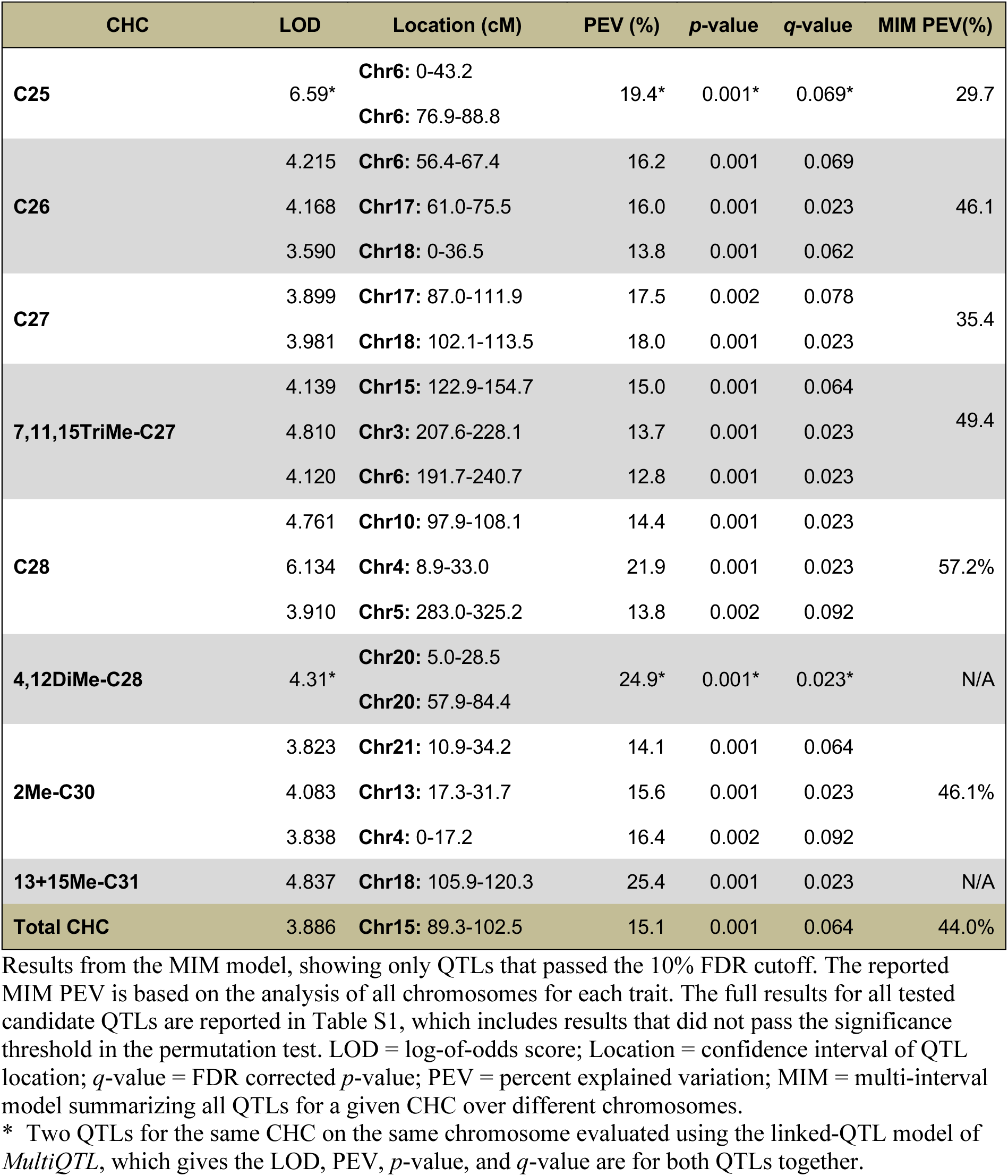
QTLs underlying variation in CHCs.

Interestingly, we did not detect statistically significant QTLs for most CHCs, but when we did find QTLs for a certain CHC we found more than one (Supp. Fig. S2A). Out of the eight CHCs with QTLs, four had three QTLs, five had two QTLs, on only one CHC had a single QTL, which would be unlikely if all CHCs had equal probability to have QTLs (Poisson dispersion test *p* = 0.003). These results indicate that the variation in some CHCs has a greater genetic component than others. QTLs with large effect size are more likely to be discovered with the limited statistical power given our sample size. Thus, other CHCs may still be affected by QTLs with smaller effect sizes, which are below our detection threshold. The portion of the phenotypic variation explained by QTLs (percent explained variation; PEV) for each CHC ranged from 29.7% to 57.2% (Table 2). The PEV for single QTLs varied between 12.8% and 25.4%. The most genetically determined CHC is C28, for which three QTLs had PEV of 14.4%, 13.8%, and 21.9%, and together accounted for 57.2% PEV, which implies that this trait is affected by genetic factors more than by environmental factors.

The 20 QTLs are located in 11 of the 26 chromosomes of *C. niger*, and the number of QTLs per chromosome was not evenly distributed (Supp. Fig. S2B; between 1-5 QTLs per chromosome; Poisson dispersion test *p* = 0.06). In some chromosomes the QTLs for different CHCs appear to be clustered and even overlapping (Fig. 3). For example, QTLs for C28 and 2Me-C30 were found together in the tellomeric region of chromosome 4. Chromosome 18 harbors one locus associated with C26 and another associated with C27, 13Me- and 15Me-C31 (the LOD plots are virtually overlapping). Each of the QTLs for C26 and C27 on chromosome 17 are also weakly associated with the other CHC. These results suggest that multiple CHCs may be regulated by the same gene or by different genes in a small genomic region, which could harbor a cluster of co-regulated CHC genes.

### Candidate genes

Our sample size (*n*=81) limited the resolution of QTL mapping, resulting in 95% confidence intervals of genetic map length ranging between 10 and 49cM, and physical sizes ranging between 345 and 3,255 Kb. We note that three of the 20 QTLs included contigs that were mapped based on the conservation of synteny with *Formica* (150 out of 565 kbp of the QTL for C25 on chromosome 6; 167 / 3,571 kbp of 4,12DiMe-C28 on chr. 9; and 101 / 1,287 kbp of C26 on chr. 18). The mapping of these contigs was confirmed by the conservation of microsynteny (gene order) with orthologous genes in *Formica*. Each interval contained between 21 and 156 putative protein coding genes. Using our data, it is impossible to narrow down these gene lists and identify the loci functionally responsible for the variation in the phenotype. Nevertheless, we noted several candidate genes in these intervals that were implicated in molecular functions and biological processes related to the synthesis of CHCs or to the biology and development of the cuticle (Table 3). For example, these include a fatty acyl-CoA reductase gene (Cnig_gene3669) in the QTL for C27 on chromosome 17 and a long-chain fatty acid elongase gene (Cnig_gene6578) in the QTL for C26 on chromosome 18.

**Table 3:** Candidate genes in QTLs underlying variation in CHCs.

| QTL | Gene | Ortholog | Description |
| --- | --- | --- | --- |
| 2Me-C30 chr. 13 | Cnig_gene5601 | FABP9_CAEEL | Fatty acid-binding protein homolog 9 |
| 2Me-C30 chr. 21 | Cnig_gene2019 | LIPI_HUMAN | Lipase member I |
| 7,11,15TriMe-C27 chr. 06 | Cnig_gene13278 | PEX2_RAT | Peroxisome biogenesis factor 2 |
| 7,11,15TriMe-C27 chr. 15 | Cnig_gene4696 | ACADV_HUMAN | Very long-chain specific acyl-CoA dehydrogenase, mitochondrial (ECO:0000305\|PubMed:7668252) |
| 7,11,15TriMe-C27 chr. 15 | Cnig_gene4695 | KAPR1_DROME | cAMP-dependent protein kinase type I regulatory subunit |
| C26 chr. 18 | Cnig_gene6578 | ELO3_CAEEL | Putative fatty acid elongation protein 3 |
| C27 chr. 17 | Cnig_gene3669 | WAT_DROME | Fatty acyl-CoA reductase *wat* (ECO:0000305) |
| Total chr. 15 | Cnig_gene4381 | FABD_DROME | Probable malonyl-CoA-acyl carrier protein transacylase, mitochondrial |

QTL; gene ID for the *C. niger* candidate gene; gene ID of the best Swissprot blast hit and the functional description of that gene.

## Discussion

Here we applied a genomic mapping approach to test for genetic effects on CHCs, which serve as major chemical cues for recognition and other functions in social insects. We had to establish a chromosome-level reference genome assembly for the desert ant *Cataglyphis niger* as the foundation for this genomic mapping study. The long-read sequencing combined with the high-resolution linkage map resulted in draft assemblies for each of the 26 linkage groups, which is in agreement with the chromosome number in *Cataglyphis hispanica*, where one lineage has 26 chromosomes while the other has 27 (Darras, et al. 2022). The linkage map and the assembly revealed an exceptionally high estimate of a genome-wide average recombination rate of 21.3 cM/Mb. This rate is similar to the record for animals that is held by honey bees, with estimates ranging between 19-22 cM/Mb (Beye, et al. 2006; Solignac, et al. 2007; Meznar, et al. 2010; Korol and Rybnikov 2024). The total physical and genetic lengths of the *C. niger* genome are both greater than those of *Apis mellifera*: 236Mb and 5041cM vs. 187Mb and 4,114cM (Solignac, et al. 2007). For the QTL mapping analysis, we analyzed CHC profiles and genomic polymorphism in a sample of 81 haploid brothers. We used RAD sequencing to obtain a high-density genetic map for QTL mapping based on 7769 markers. This density of markers is more than needed for this progeny sample. The QTL detection and the resolution of QTL location is limited by the number of recombination events in our sample rather than the number of genetic markers. We report associations between 19 QTLs and 8 out of 31 CHCs tested, and one locus that is associated with the total amount of CHCs. We identified rather large effect sizes that each explain between 13% and 25% of the variation in a given CHC. Between 30% and 57% of the variation of the eight affected CHCs was explained by the combined genetic effects. Our study opens a window to the genomic architecture underlying CHC variation in a social insect species.

Our results add to a large volume of literature on determinants of CHC profiles. Previous studies demonstrated large environmental effects by various factors, including diet (Liang and Silverman 2000), pathogens (Tranter, et al. 2015; Leclerc and Detrain 2016), and abiotic environmental factors such as temperature and humidity (Silva, et al. 2024). Incorporating our results allows a comprehensive understanding of the mechanisms shaping CHC profiles by a combination of genetic and environmental effects. Our results reveal large variation in the genetic effects on different CHCs. We discovered QTLs with intermediate effect sizes for eight CHCs; each locus explaining between 12 and 25 percent of phenotypic variation. Up to 57% of the variation of C28 was explained by the combined effects of the QTLs we identified, but for the other CHCs most of the phenotypic variation was not explained by the genetic effects. CHCs with QTLs are expected to be heritable as suggested by previous studies in multiple ant species (Nowbahari, et al. 1990; Lahav, et al. 1999; Akino, et al. 2004; Ozaki, et al. 2005; van Zweden, et al. 2009; Walsh, et al. 2020). Our results are in line with estimates of narrow-sense heritability (*h^2^*) of individual CHCs in the ant *Monomorium pharaonis* that ranged between 0 and 0.36 (Walsh, et al. 2020). We could not detect any QTLs associated with 23 out of the 31 CHCs in our data. This may be because some CHCs are only affected by small effect size QTLs that could not be detected with our sample size. Generally, our estimates of the total genetic component for any of the studied CHCs should be considered as a lower bound, because they may be affected by additional small effect size QTLs that we could not detect. Nevertheless, our results indicate substantial variation among CHCs in the magnitude of their genetic basis and that for most CHCs the genetic component is smaller than the environmental component. Environmental effects may dominate the cues for nestmate recognition because nestmates share the same abiotic environment of the nest, the same diet, and other biotic effects such as pathogens.

The eight CHCs with associated QTLs include four straight-chain and four methyl-branched alkanes. This may be surprising considering that methyl-branched CHCs have greater information content, and thus are expected to be under greater selection as recognition cues, and more likely to evolve a genetic basis. This hypothesis was supported by some studies that reported stronger differentiation between colonies by methyl-branched vs. straight alkanes in some species (e.g., in *Cataglyphis iberica*; Dahbi, et al. 1996) and stronger direct and indirect genetic effects (e.g., in *Formica rufibarbis*; Van Zweden, et al. 2010). There may be other reasons for the evolution of a genetic basis for the less informative straight-chain alkanes, other than for nestmate recognition purposes. For example, these alkanes may be used for simple, conserved signals that do not require high information content, such as differentiating between foragers and nurses (Greene and Gordon 2003; Martin and Drijfhout 2009b) or to identify a mated queen (Hora, et al. 2008). If alkanes are used for such a conserved function they may evolve a genetic basis to better regulate their expression.

As discussed in the Introduction, the most obvious candidate genes are ant homologs of enzymes that were extensively studied in model insects and implicated in the biosynthetic pathway of long-chain fatty acids, which are the precursors of CHCs. These include fatty acid synthases, fatty acid elongases, and fatty acid reductases. One such candidate, Cnig_gene3669 in the QTL for C27 on chromosome 17, is a homolog of *Drosophila* fatty acyl-CoA reductase (FAR) genes. The top blast hit in *Drosophila* is the gene *waterproof* (*wat*; see a gene tree of ant and fly FAR genes in Supp. Fig. S3), which is involved in tracheal development. Knocking out *wat* interfered with gas filling of the tracheal tubes during *Drosophila* embryogenesis (Jaspers, et al. 2014) and *wat* homologs in planthoppers affect CHC quantities (Li, et al. 2019; Li, et al. 2020). Generally, FAR genes are responsible for one of the last steps in CHC biosynthetic pathways, where fatty-acyl CoA precursors are converted to hydrocarbons. The FAR gene family was implicated in CHC synthesis in *Drosophila* based on their expression in oenocytes and their role in regulation of oenocyte development (Finet, et al. 2019), association with intraspecific CHC variation, and knockdown experiments that affected CHC length (Dembeck, et al. 2015). Another candidate, Cnig_gene6578 in the QTL for C26 on chromosome 18, is a homolog of *Drosophila* long-chain fatty acid elongase genes. This gene family takes part in the CHC synthesis pathway, in elongating the chain length of fatty acids initially synthesized by fatty acid synthases beyond 20 carbons (Blomquist and Bagneres 2010). Knockdown of such elongases in *Drosophila* and in the brown planthopper *Nilaparvata lugens* affected abundances of different CHCs (Holze, et al. 2021). The QTL for 2Me-C30 on chromosome 21 contains a homolog of human lipase member I, which hydrolyzes phosphatidic acid (PA) to 2-acyl lysophosphatidic acid (LPA; a potent bioactive lipid mediator) and a fatty acid (Hiramatsu, et al. 2003). While the function of the insect homolog is unknown, it is a promising candidate in lipid metabolism, which may be involved in the synthesis of CHC precursors.

In this study we used a cohort of male brothers as the mapping population. Since this cohort was collected from a single nest, it represent a sample of a single colony from the population. As such, we cannot see how the genetic architecture of CHCs varies across the population. Nevertheless, given the strong evidence for multiple QTLs for multiple CHCs in this colony sample we expect this to generally characterize the genetic architecture of ant CHCs, even if the specific loci and their effect sizes may vary. Our motivation for using males was that they are haploid, which is advantageous for genomic mapping, and because they developed in the same nest in uniform environmental conditions. Male CHCs are likely to have functional roles for recognition and discrimination in mate choice during mating flights. CHCs were extensively investigated in such roles in non-social insects. For example, male offspring from a cross between the wasps *Nasonia vitripennis* and *N. giraulti* were used for a QTL mapping study of CHCs (Niehuis, et al. 2011). CHCs were demonstrated to function as cues for mate choice in this system, which allows females to discriminate between males from different species (Buellesbach, et al. 2013). Similarly, CHCs of male ants are likely to function as cues for mate choice by queens (Sprenger and Menzel 2020). In the worker caste of social insects, CHCs are most consequential in their role as nestmate recognition cues for identifying intruders from other colonies (van Zweden and d’Ettorre 2010). The genetic effects we uncovered in males are likely to also affect CHC profiles of queens and workers, because most CHCs are observed in more than one caste of the same species (Hefetz 2007). Genes involved in the synthesis of these CHCs likely evolved in response to selection pressures on a range of various functional roles in each of the different castes (e.g. nestmate recognition in workers vs. mate choice in males). Natural selection on the ability to discriminate against foreign workers may increase genetic polymorphism that contributes to CHC variation, and this polymorphism may also affect variation of the same CHC in males even if this CHC is not under selection in males. Therefore, some of the QTLs we detected in males may actually be functionally important in one or the other caste. Further studies in other castes will provide a more comprehensive understanding of this genetic system and its functional significance.

## Acknowledgements

We thank Abraham Hefetz for advice and access to equipment for chemical analysis of CHCs. We thank Jessica Purcell and Alan Brelsford for advice on RAD sequencing. This study was funded by Israel Science Foundation grant number 646/15.

## Data availability

All sequence data was deposited to NCBI: the new genome assembly Cnig_gn3.1 is under NCBI BioProject accession PRJNA1168153 and the raw Pacbio sequencing reads are under SRA accession SRR10302333. The RAD-seq data is under Bioproject accession PRJNA1000554. Chemical data is provided as supplementary materials to this paper.

## Supplementary

**Supplementary Figure S1:**
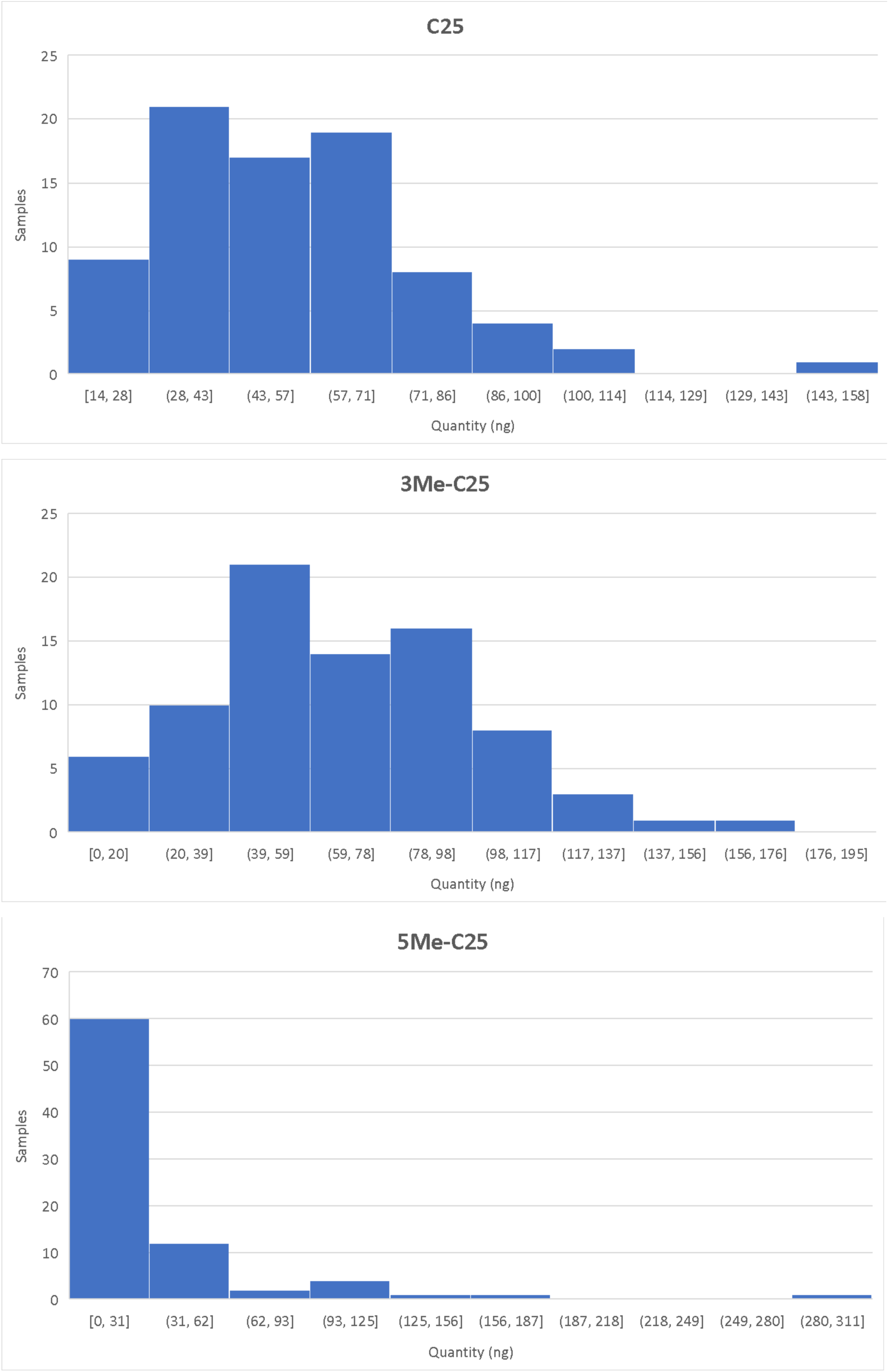
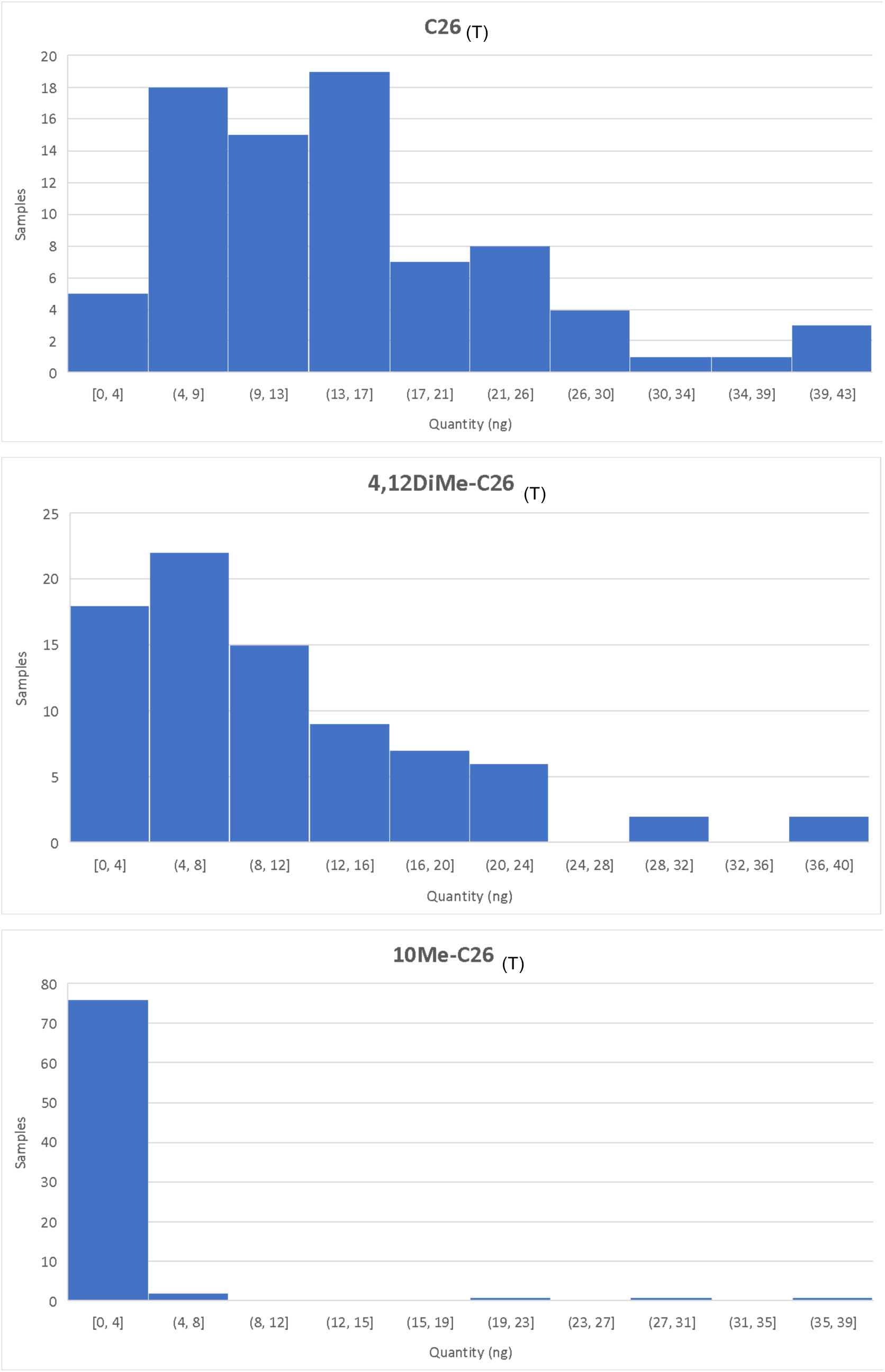
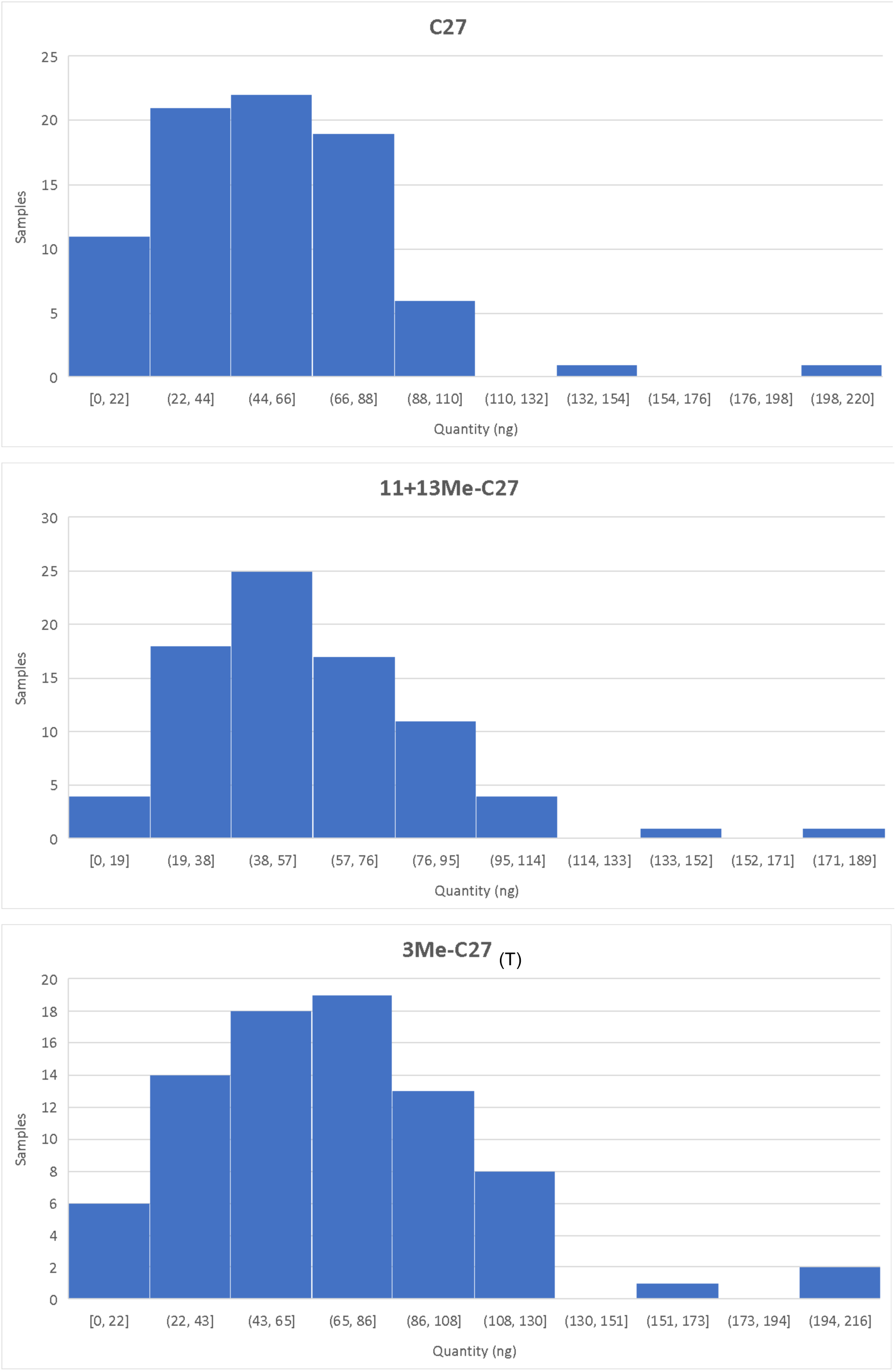
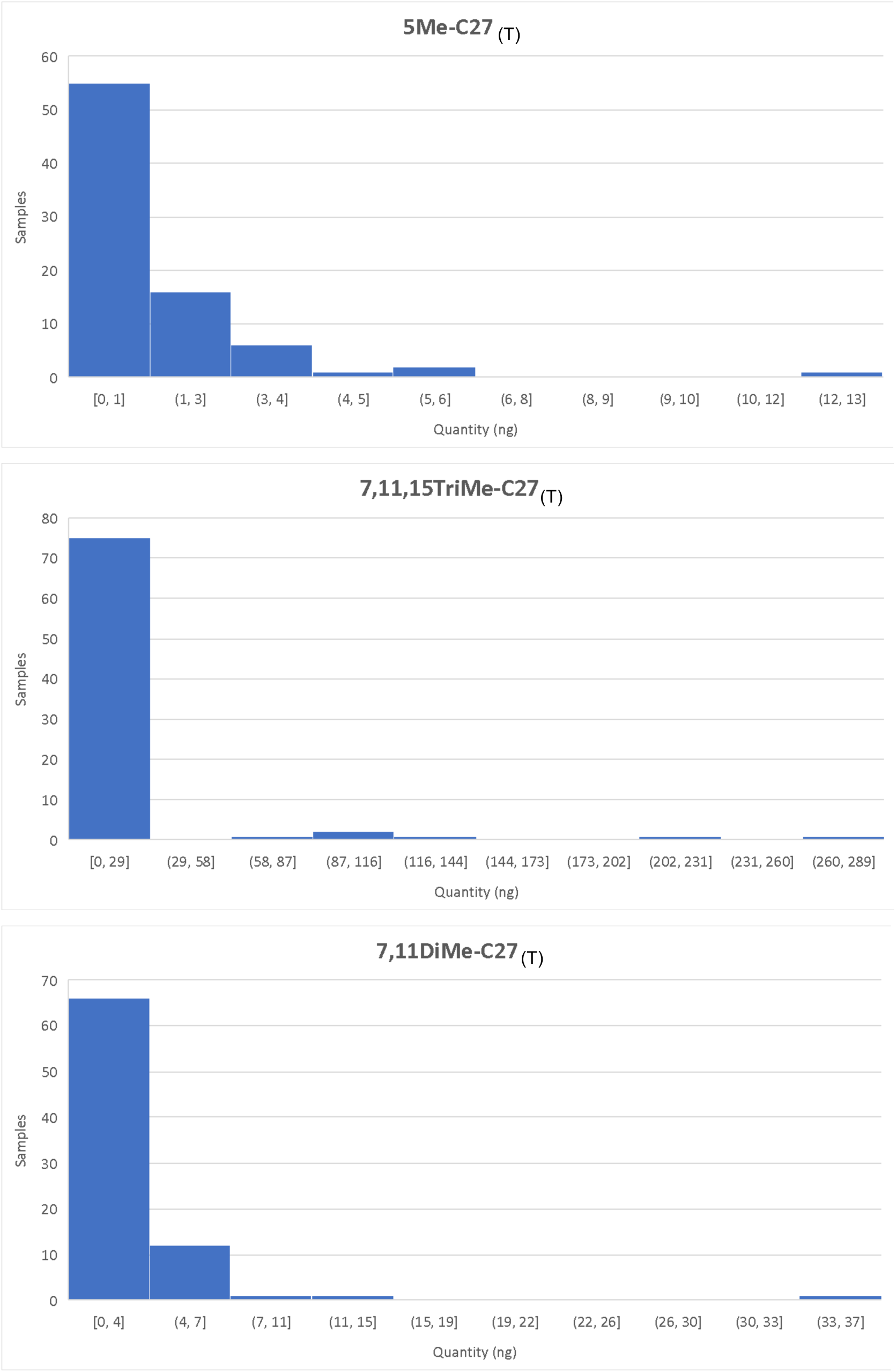
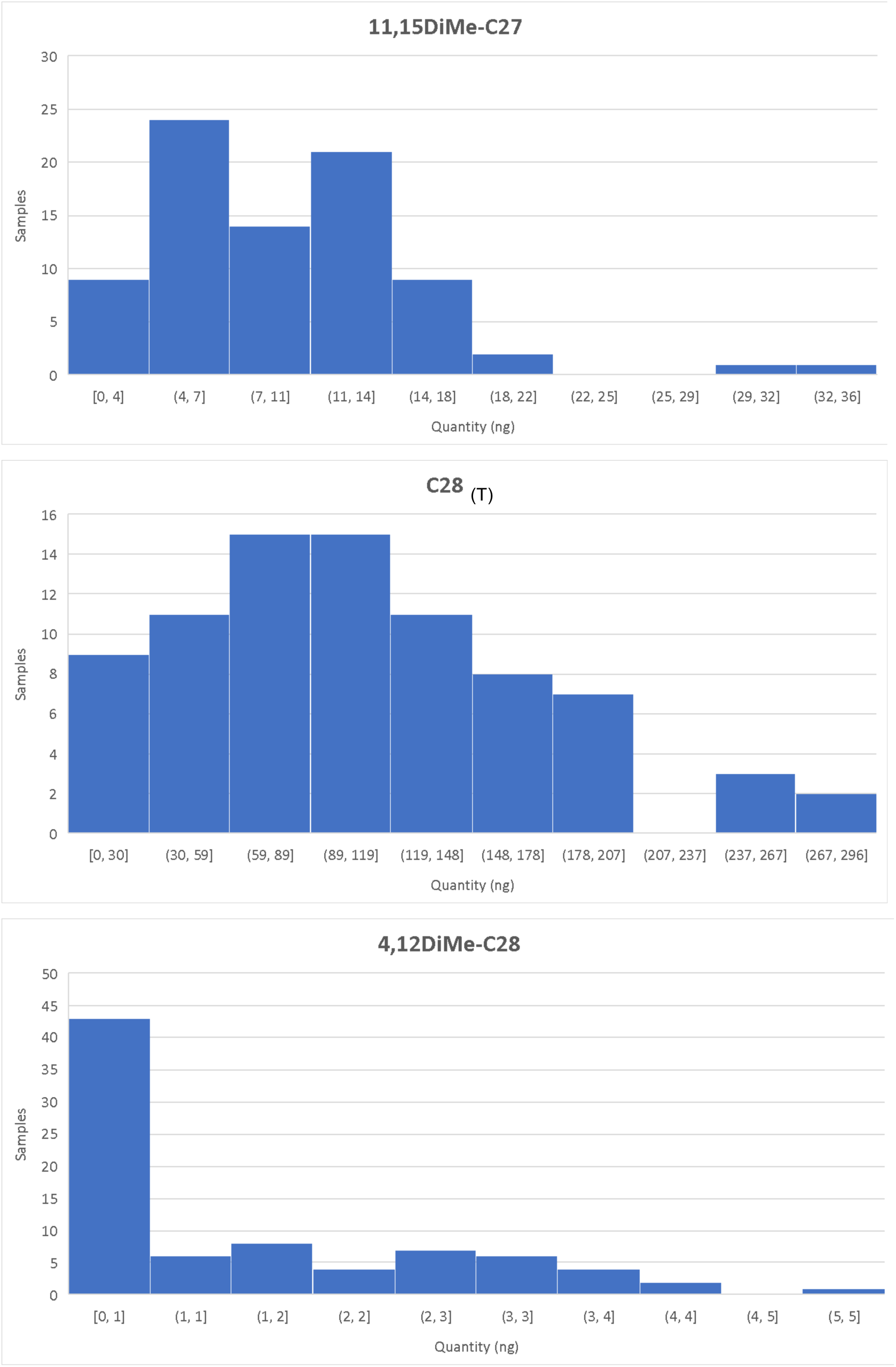
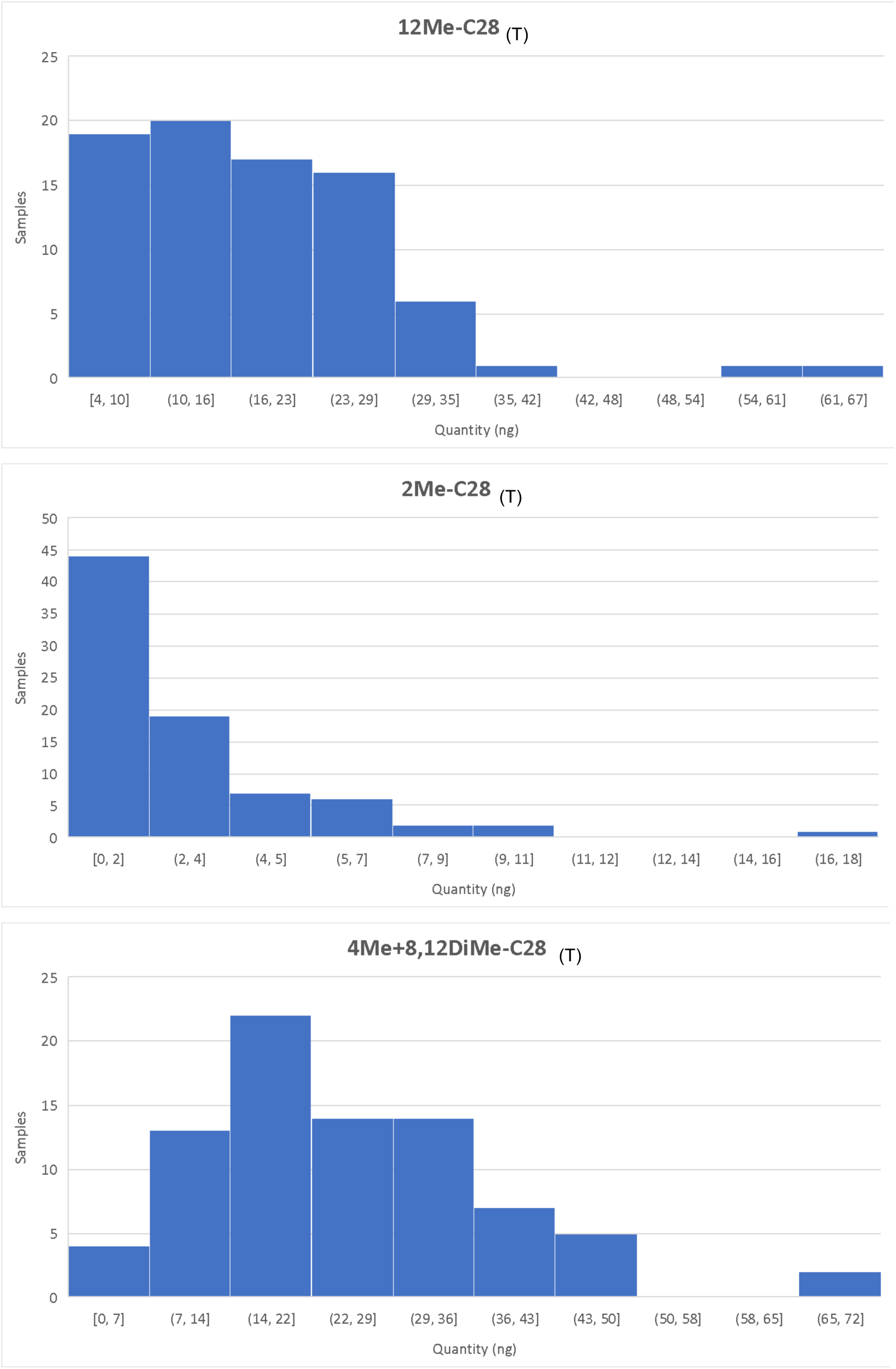
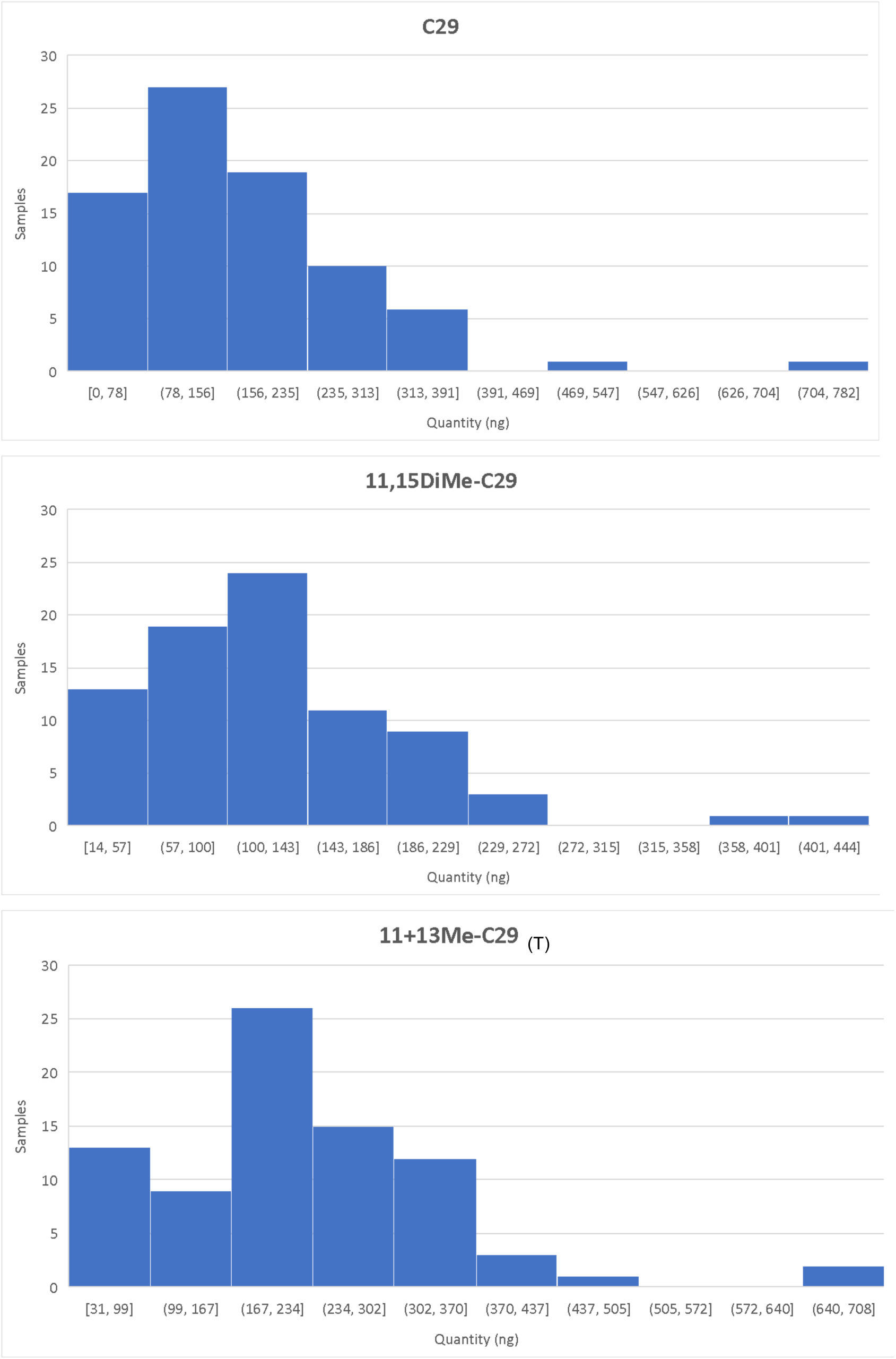
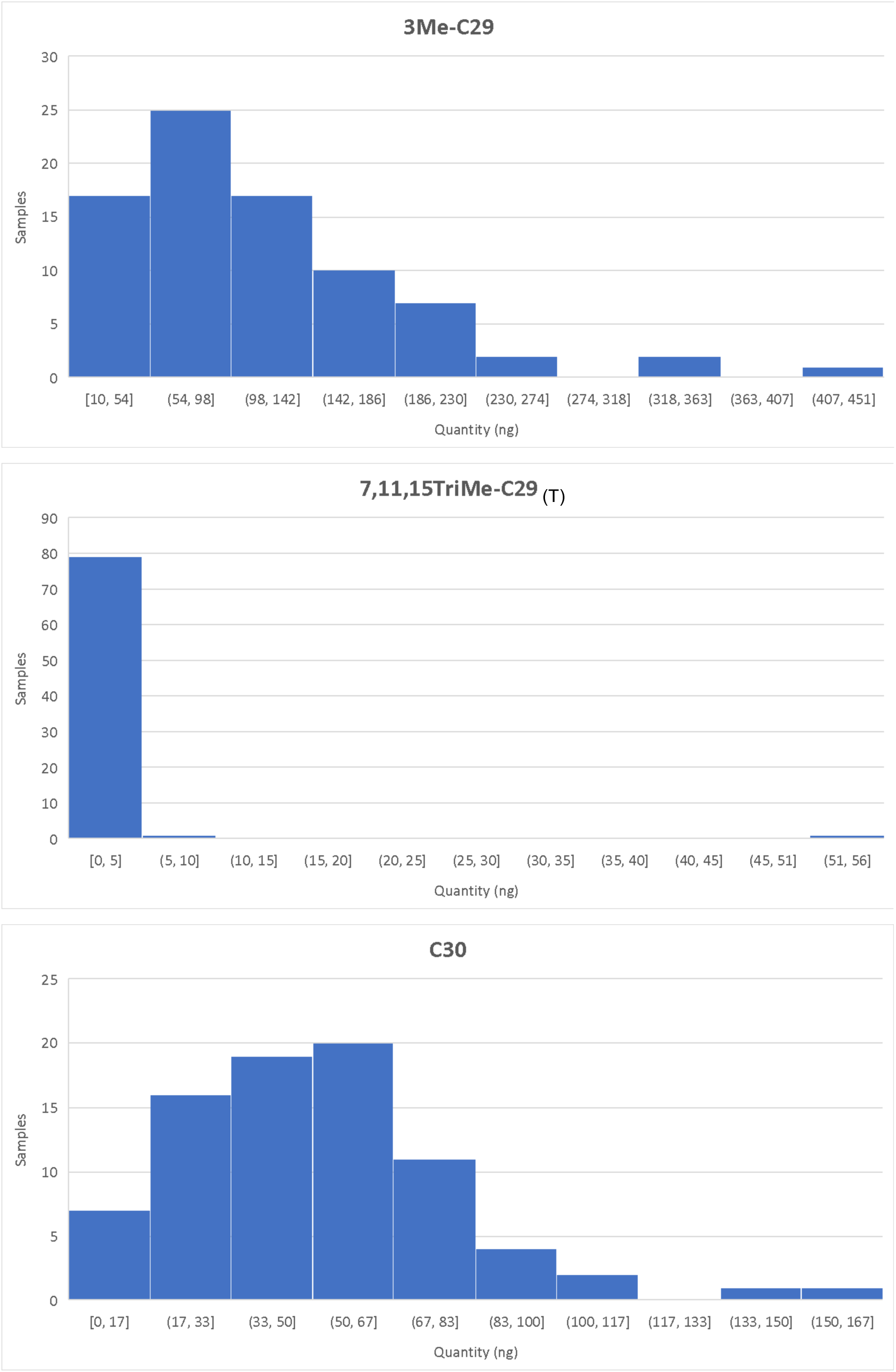
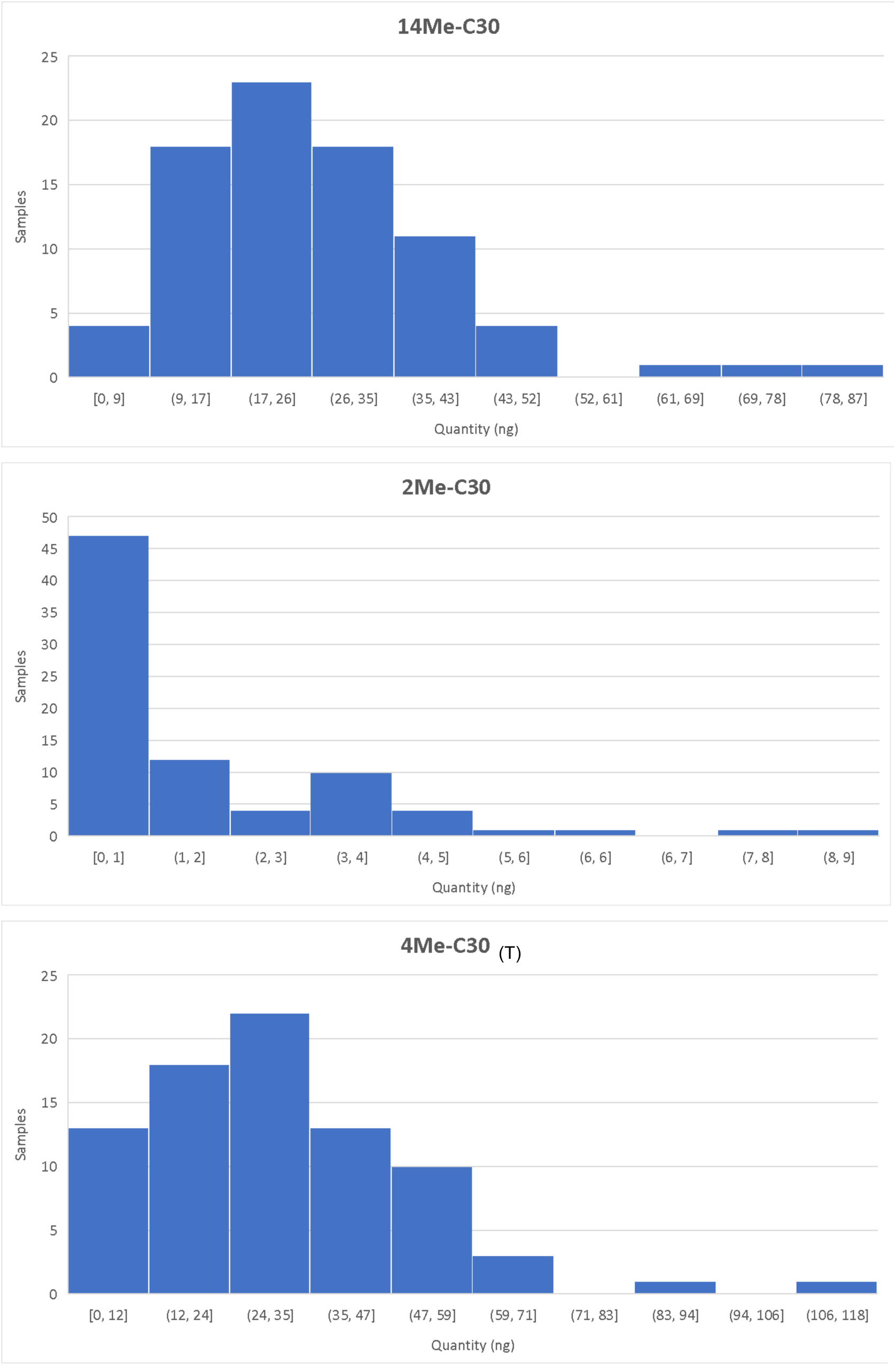
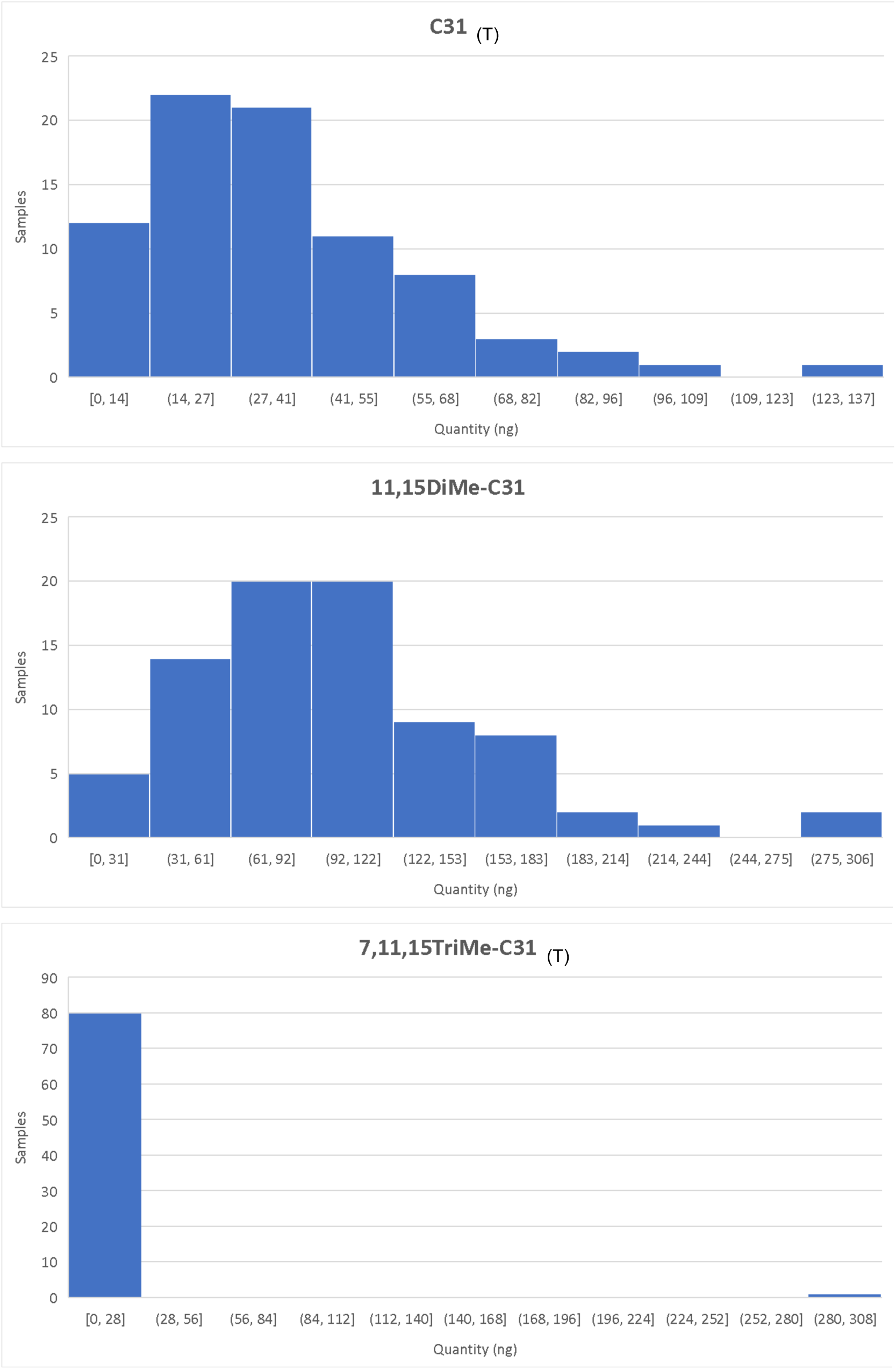
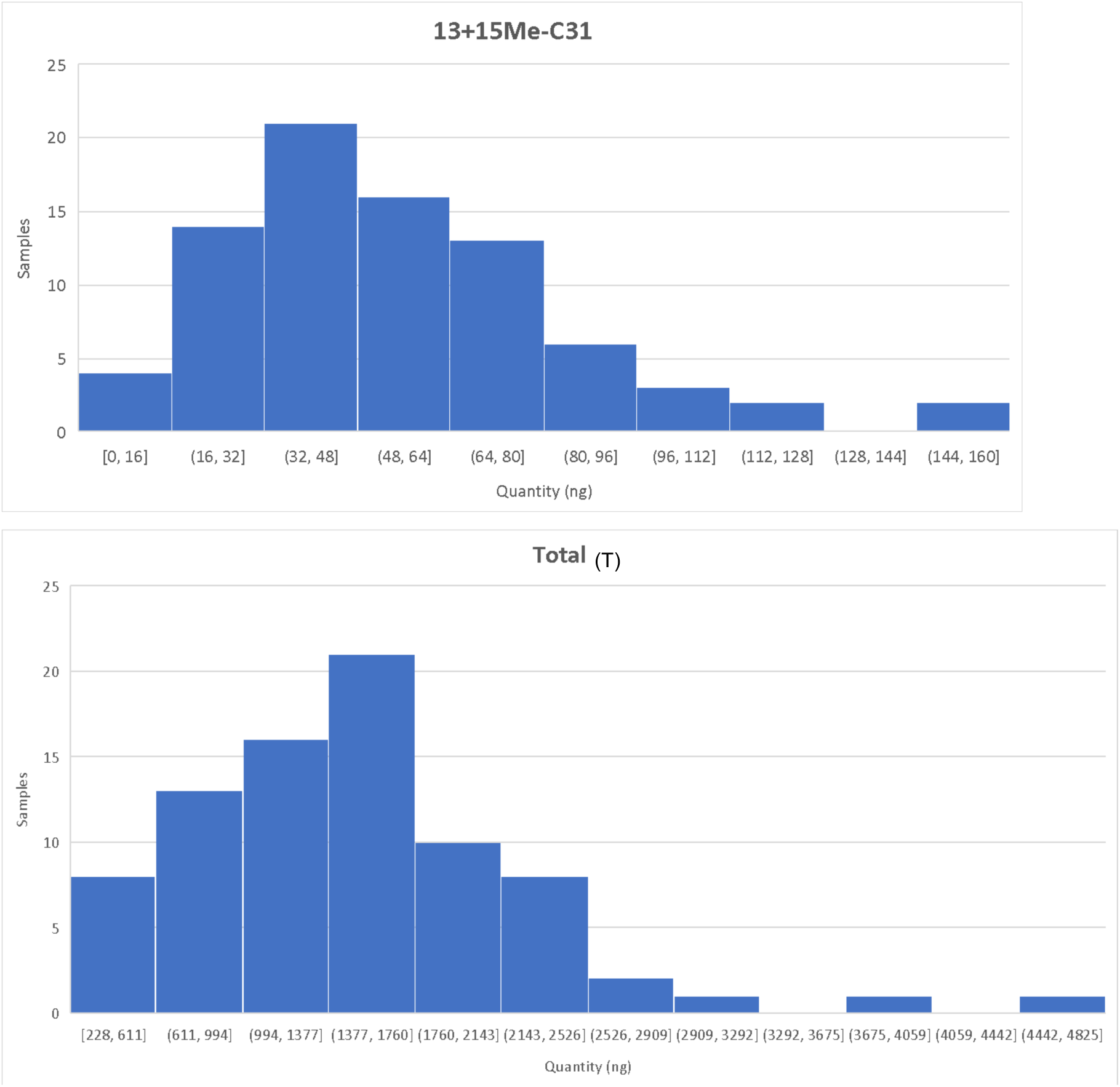
Distributions of the absolute quantity of each CHC in the samples. CHCs marked with (T) were log-transformed.

**Supplementary Figure S2:**
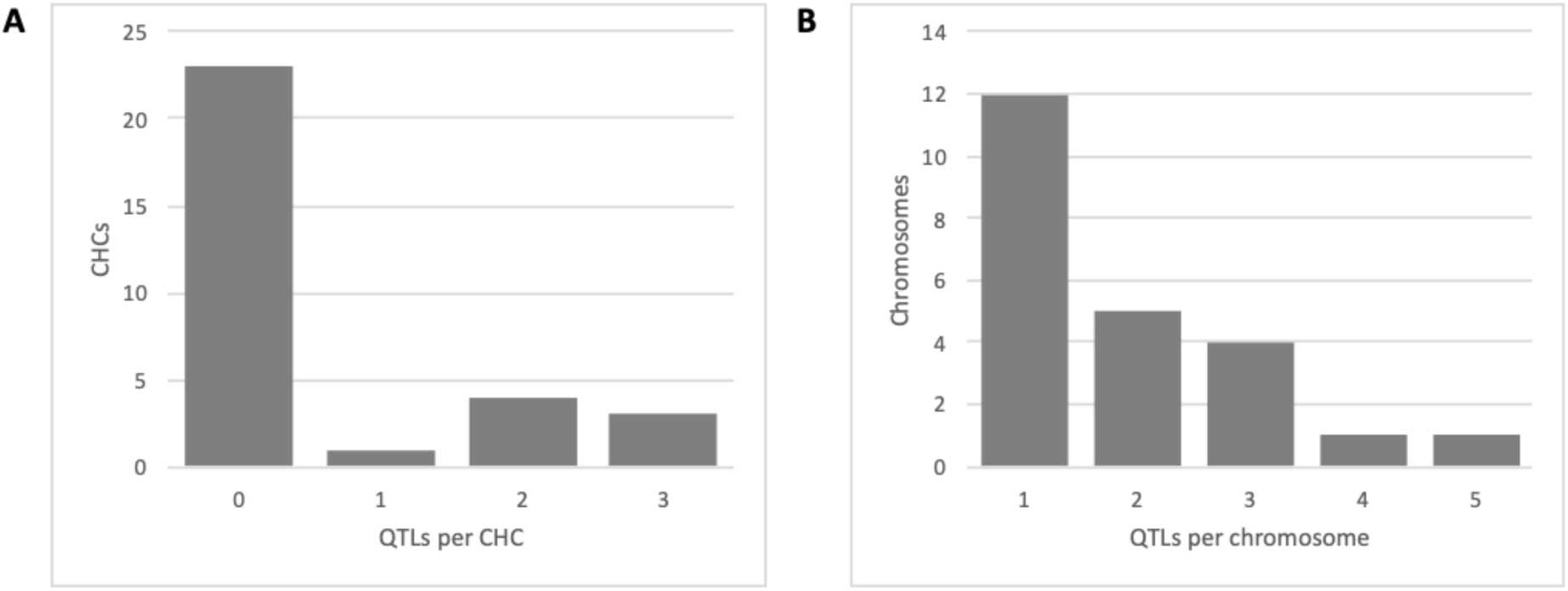
Distribution of the number of QTLs per CHC (A) and QTLs per chromosome (B).

**Supplementary Figure S3:**
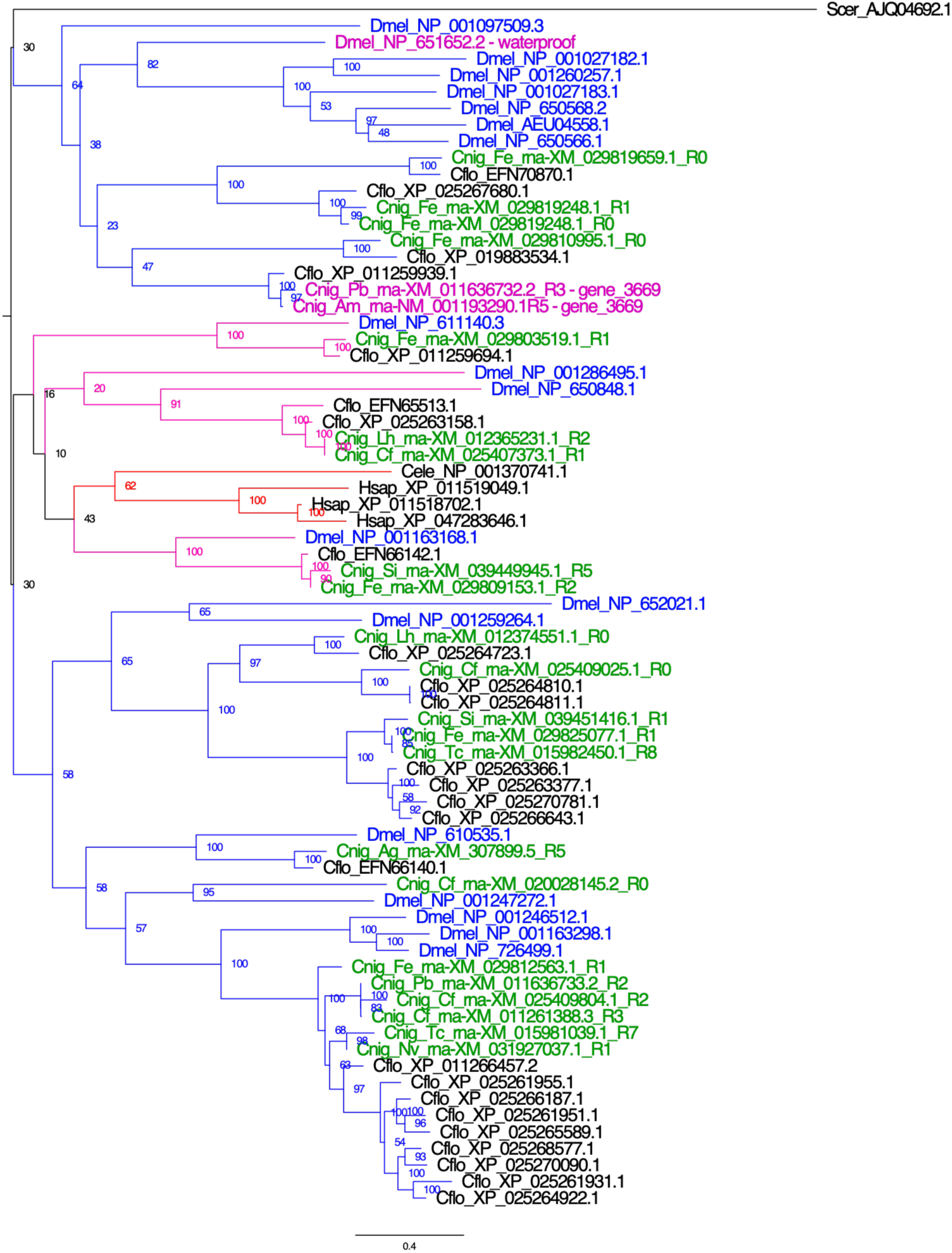
Fatty acyl-CoA reductase gene tree. Including fly (*Drosophila melanogaster*) and ant (*Cataglyphis niger* and *Camponotus floridanus*) genes. The *Drosophila waterproof* gene and other fly paralogs are the closest homologs in *Drosophila* to *C. niger* gene 3669 in the QTL for C27. The gene names are colored by species and subfamilies that existed in the common ancestor of ants and flies are colored in distinct colors.

**Supplementary Table S1:**
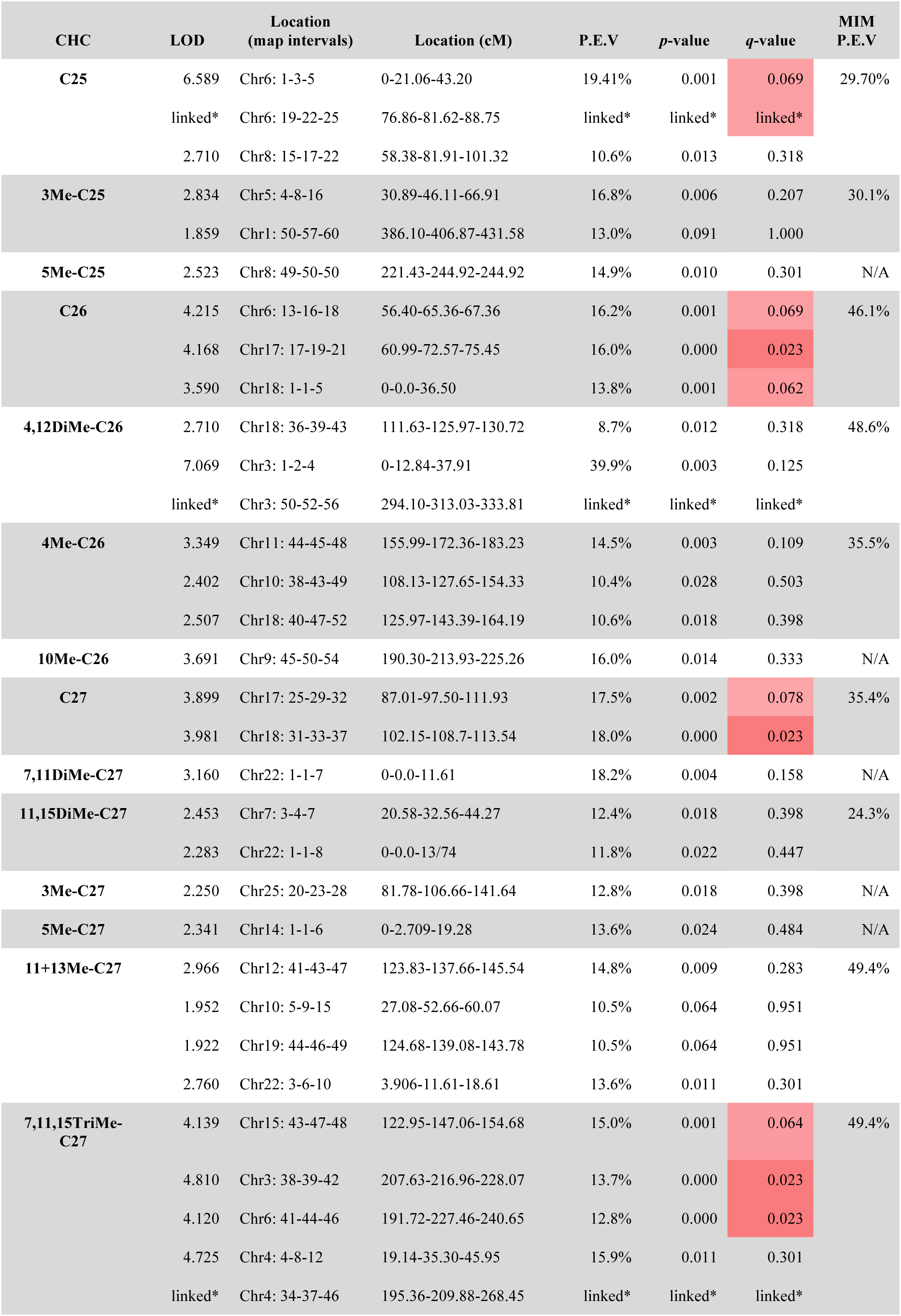
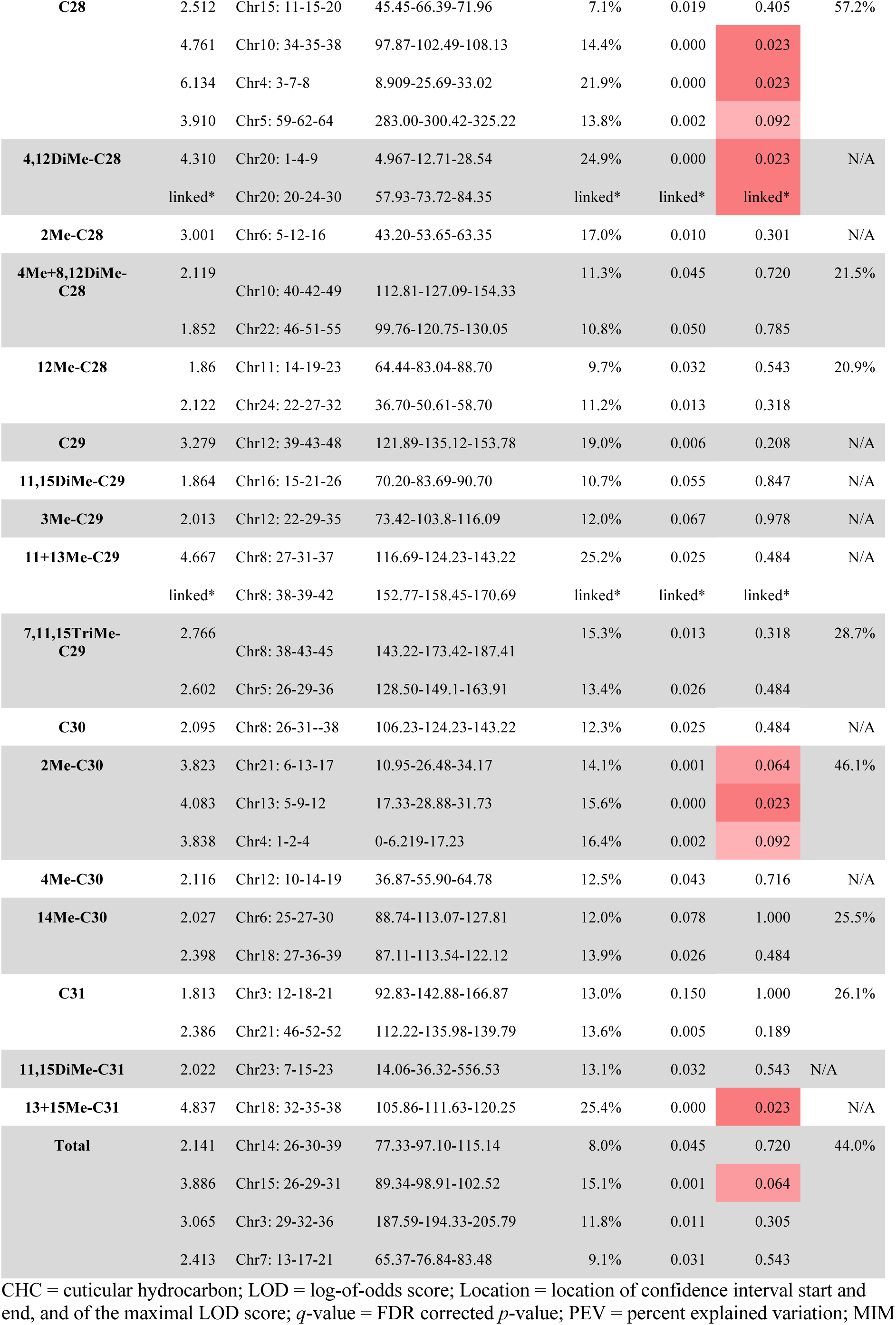
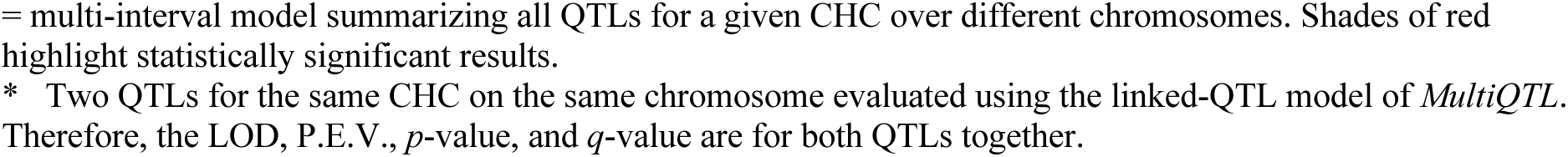
QTLs underlying variation in CHCs (full table of tested candidate QTLs, including non-significant results)

| CHC | LOD | Location<br>(map intervals) | Location (cM) | P.E.V | *p*-value | *q*-value | MIM<br>P.E.V |
| --- | --- | --- | --- | --- | --- | --- | --- |
| **C25** | 6.589 | Chr6: 1-3-5 | 0-21.06-43.20 | 19.41% | 0.001 | 0.069 | 29.70% |
|  | linked* | Chr6: 19-22-25 | 76.86-81.62-88.75 | linked* | linked* | linked* |  |
|  | 2.710 | Chr8: 15-17-22 | 58.38-81.91-101.32 | 10.6% | 0.013 | 0.318 |  |
| **3Me-C25** | 2.834 | Chr5: 4-8-16 | 30.89-46.11-66.91 | 16.8% | 0.006 | 0.207 | 30.1% |
|  | 1.859 | Chr1: 50-57-60 | 386.10-406.87-431.58 | 13.0% | 0.091 | 1.000 |  |
| **5Me-C25** | 2.523 | Chr8: 49-50-50 | 221.43-244.92-244.92 | 14.9% | 0.010 | 0.301 | N/A |
| **C26** | 4.215 | Chr6: 13-16-18 | 56.40-65.36-67.36 | 16.2% | 0.001 | 0.069 | 46.1% |
|  | 4.168 | Chr17: 17-19-21 | 60.99-72.57-75.45 | 16.0% | 0.000 | 0.023 |  |
|  | 3.590 | Chr18: 1-1-5 | 0-0.0-36.50 | 13.8% | 0.001 | 0.062 |  |
| **4,12DiMe-C26** | 2.710 | Chr18: 36-39-43 | 111.63-125.97-130.72 | 8.7% | 0.012 | 0.318 | 48.6% |
|  | 7.069 | Chr3: 1-2-4 | 0-12.84-37.91 | 39.9% | 0.003 | 0.125 |  |
|  | linked* | Chr3: 50-52-56 | 294.10-313.03-333.81 | linked* | linked* | linked* |  |
| **4Me-C26** | 3.349 | Chr11: 44-45-48 | 155.99-172.36-183.23 | 14.5% | 0.003 | 0.109 | 35.5% |
|  | 2.402 | Chr10: 38-43-49 | 108.13-127.65-154.33 | 10.4% | 0.028 | 0.503 |  |
|  | 2.507 | Chr18: 40-47-52 | 125.97-143.39-164.19 | 10.6% | 0.018 | 0.398 |  |
| **10Me-C26** | 3.691 | Chr9: 45-50-54 | 190.30-213.93-225.26 | 16.0% | 0.014 | 0.333 | N/A |
| **C27** | 3.899 | Chr17: 25-29-32 | 87.01-97.50-111.93 | 17.5% | 0.002 | 0.078 | 35.4% |
|  | 3.981 | Chr18: 31-33-37 | 102.15-108.7-113.54 | 18.0% | 0.000 | 0.023 |  |
| **7,11DiMe-C27** | 3.160 | Chr22: 1-1-7 | 0-0.0-11.61 | 18.2% | 0.004 | 0.158 | N/A |
| **11,15DiMe-C27** | 2.453 | Chr7: 3-4-7 | 20.58-32.56-44.27 | 12.4% | 0.018 | 0.398 | 24.3% |
|  | 2.283 | Chr22: 1-1-8 | 0-0.0-13/74 | 11.8% | 0.022 | 0.447 |  |
| **3Me-C27** | 2.250 | Chr25: 20-23-28 | 81.78-106.66-141.64 | 12.8% | 0.018 | 0.398 | N/A |
| **5Me-C27** | 2.341 | Chr14: 1-1-6 | 0-2.709-19.28 | 13.6% | 0.024 | 0.484 | N/A |
| **11+13Me-C27** | 2.966 | Chr12: 41-43-47 | 123.83-137.66-145.54 | 14.8% | 0.009 | 0.283 | 49.4% |
|  | 1.952 | Chr10: 5-9-15 | 27.08-52.66-60.07 | 10.5% | 0.064 | 0.951 |  |
|  | 1.922 | Chr19: 44-46-49 | 124.68-139.08-143.78 | 10.5% | 0.064 | 0.951 |  |
|  | 2.760 | Chr22: 3-6-10 | 3.906-11.61-18.61 | 13.6% | 0.011 | 0.301 |  |
| **7,11,15TriMe-C27** | 4.139 | Chr15: 43-47-48 | 122.95-147.06-154.68 | 15.0% | 0.001 | 0.064 | 49.4% |
|  | 4.810 | Chr3: 38-39-42 | 207.63-216.96-228.07 | 13.7% | 0.000 | 0.023 |  |
|  | 4.120 | Chr6: 41-44-46 | 191.72-227.46-240.65 | 12.8% | 0.000 | 0.023 |  |
|  | 4.725 | Chr4: 4-8-12 | 19.14-35.30-45.95 | 15.9% | 0.011 | 0.301 |  |
|  | linked* | Chr4: 34-37-46 | 195.36-209.88-268.45 | linked* | linked* | linked* |  |
| **C28** | 2.512 | Chr15: 11-15-20 | 45.45-66.39-71.96 | 7.1% | 0.019 | 0.405 | 57.2% |
|  | 4.761 | Chr10: 34-35-38 | 97.87-102.49-108.13 | 14.4% | 0.000 | 0.023 |  |
|  | 6.134 | Chr4: 3-7-8 | 8.909-25.69-33.02 | 21.9% | 0.000 | 0.023 |  |
|  | 3.910 | Chr5: 59-62-64 | 283.00-300.42-325.22 | 13.8% | 0.002 | 0.092 |  |
| **4,12DiMe-C28** | 4.310 | Chr20: 1-4-9 | 4.967-12.71-28.54 | 24.9% | 0.000 | 0.023 | N/A |
|  | linked* | Chr20: 20-24-30 | 57.93-73.72-84.35 | linked* | linked* | linked* |  |
| **2Me-C28** | 3.001 | Chr6: 5-12-16 | 43.20-53.65-63.35 | 17.0% | 0.010 | 0.301 | N/A |
| **4Me+8,12DiMe-C28** | 2.119 | Chr10: 40-42-49 | 112.81-127.09-154.33 | 11.3% | 0.045 | 0.720 | 21.5% |
|  | 1.852 | Chr22: 46-51-55 | 99.76-120.75-130.05 | 10.8% | 0.050 | 0.785 |  |
| **12Me-C28** | 1.86 | Chr11: 14-19-23 | 64.44-83.04-88.70 | 9.7% | 0.032 | 0.543 | 20.9% |
|  | 2.122 | Chr24: 22-27-32 | 36.70-50.61-58.70 | 11.2% | 0.013 | 0.318 |  |
| **C29** | 3.279 | Chr12: 39-43-48 | 121.89-135.12-153.78 | 19.0% | 0.006 | 0.208 | N/A |
| **11,15DiMe-C29** | 1.864 | Chr16: 15-21-26 | 70.20-83.69-90.70 | 10.7% | 0.055 | 0.847 | N/A |
| **3Me-C29** | 2.013 | Chr12: 22-29-35 | 73.42-103.8-116.09 | 12.0% | 0.067 | 0.978 | N/A |
| **11+13Me-C29** | 4.667 | Chr8: 27-31-37 | 116.69-124.23-143.22 | 25.2% | 0.025 | 0.484 | N/A |
|  | linked* | Chr8: 38-39-42 | 152.77-158.45-170.69 | linked* | linked* | linked* |  |
| **7,11,15TriMe-C29** | 2.766 | Chr8: 38-43-45 | 143.22-173.42-187.41 | 15.3% | 0.013 | 0.318 | 28.7% |
|  | 2.602 | Chr5: 26-29-36 | 128.50-149.1-163.91 | 13.4% | 0.026 | 0.484 |  |
| **C30** | 2.095 | Chr8: 26-31--38 | 106.23-124.23-143.22 | 12.3% | 0.025 | 0.484 | N/A |
| **2Me-C30** | 3.823 | Chr21: 6-13-17 | 10.95-26.48-34.17 | 14.1% | 0.001 | 0.064 | 46.1% |
|  | 4.083 | Chr13: 5-9-12 | 17.33-28.88-31.73 | 15.6% | 0.000 | 0.023 |  |
|  | 3.838 | Chr4: 1-2-4 | 0-6.219-17.23 | 16.4% | 0.002 | 0.092 |  |
| **4Me-C30** | 2.116 | Chr12: 10-14-19 | 36.87-55.90-64.78 | 12.5% | 0.043 | 0.716 | N/A |
| **14Me-C30** | 2.027 | Chr6: 25-27-30 | 88.74-113.07-127.81 | 12.0% | 0.078 | 1.000 | 25.5% |
|  | 2.398 | Chr18: 27-36-39 | 87.11-113.54-122.12 | 13.9% | 0.026 | 0.484 |  |
| **C31** | 1.813 | Chr3: 12-18-21 | 92.83-142.88-166.87 | 13.0% | 0.150 | 1.000 | 26.1% |
|  | 2.386 | Chr21: 46-52-52 | 112.22-135.98-139.79 | 13.6% | 0.005 | 0.189 |  |
| **11,15DiMe-C31** | 2.022 | Chr23: 7-15-23 | 14.06-36.32-556.53 | 13.1% | 0.032 | 0.543 | N/A |
| **13+15Me-C31** | 4.837 | Chr18: 32-35-38 | 105.86-111.63-120.25 | 25.4% | 0.000 | 0.023 | N/A |
| **Total** | 2.141 | Chr14: 26-30-39 | 77.33-97.10-115.14 | 8.0% | 0.045 | 0.720 | 44.0% |
|  | 3.886 | Chr15: 26-29-31 | 89.34-98.91-102.52 | 15.1% | 0.001 | 0.064 |  |
|  | 3.065 | Chr3: 29-32-36 | 187.59-194.33-205.79 | 11.8% | 0.011 | 0.305 |  |
|  | 2.413 | Chr7: 13-17-21 | 65.37-76.84-83.48 | 9.1% | 0.031 | 0.543 |  |

CHC = cuticular hydrocarbon; LOD = log-of-odds score; Location = location of confidence interval start and end, and of the maximal LOD score; *q*-value = FDR corrected *p*-value; PEV = percent explained variation; MIM
= multi-interval model summarizing all QTLs for a given CHC over different chromosomes. Shades of red highlight statistically significant results. \* Two QTLs for the same CHC on the same chromosome evaluated using the linked-QTL model of *MultiQTL*. Therefore, the LOD, P.E.V., $p$ -value, and $q$ -value are for both QTLs together.

